# SOFIevaluator: a strategy for the quantitative quality assessment of SOFI data

**DOI:** 10.1101/802199

**Authors:** Benjamien Moeyaert, Wim Vandenberg, Peter Dedecker

## Abstract

Super-resolution fluorescence imaging techniques allow optical imaging of specimens beyond the diffraction limit of light. Super-resolution optical fluctuation imaging (SOFI) relies on computational analysis of stochastic blinking events to obtain a super-resolved image. As with some other super-resolution methods, this strong dependency on computational analysis can make it difficult to gauge how well the resulting images reflect the underlying sample structure. We herein report SOFIevaluator, an unbiased and parameter-free algorithm for calculating a set of metrics that describes the quality of super-resolution fluorescence imaging data for SOFI. We additionally demonstrate how SOFIevaluator can be used to identify fluorescent proteins that perform well for SOFI imaging under different imaging conditions.

## 1. Introduction

The diffraction of light has long limited light-based microscopy to a spatial resolution of approximately 200 nm in the lateral direction. The recent introduction of numerous types of sub-diffraction or super-resolution fluorescence microscopy, however, has opened new opportunities for optical imaging with diffraction-unlimited resolution in a way that is highly compatible with biological samples. Super-resolution optical fluctuation imaging (SOFI) [1] is a super-resolution fluorescence imaging approach that uses fluctuations in fluorescence dye emission to calculate an image that has a resolution that is better than the diffraction limit. Its main distinguishing feature is that it is relatively insensitive to background signal, and can be performed over a broad range of labelling densities and probe brightness [2, 3]. This robustness has also made it possible to use the technique for the first sub-diffraction observation of kinase activity [4].

However, as is often the case with super-resolution fluorescence imaging techniques, SOFI images are generated by computational processing of datasets containing hundreds or thousands of images [5–8]. The large datasets and automated processing that is required, make it difficult to establish an accurate and unbiased feel for the imaging reliability, casting uncertainty on whether a particular interpretation of the images is sustained by the data. In addition, even with perfect imaging, the available information may be limited, as is the case with low labeling densities. Traditionally, such issues have been tackled by repeating the measurement several times, but true repetitive imaging is difficult to achieve in the face of issues such as biological heterogeneity, photobleaching and phototoxicity. Several computational techniques for handling this uncertainty have been described in literature [9–15]. However, these techniques are either specific to other approaches, or are otherwise more generic and therefore not able to take advantage of direct knowledge of the SOFI imaging process. For instance, we have tried applying Fourier ring correlation to SOFI data [16], but found that in non-ideal circumstances, the apparent resolution resulting from these calculations were not consistent with the input data. These inconsistencies were most likely derived from the sample structure and camera imperfections.

We previously developed a strategy for the model-free assessment of the measurement uncertainty associated with every pixel in the SOFI imaging, by using statistical resampling of the experimental data [17]. However, this work did not provide any information on the source of these uncertainties, which can be affected by measurement noise, bleaching of the fluorophores, or imperfections in the detector. In this work we expand this work by developing a strategy to assess the overall quality of SOFI images, which we call ‘SOFIevaluator’. The SOFIevaluator is an algorithm that determines the signal-to-noise ratio of the images, calculates the fraction of SOFI signal that is due to bleaching, determines the effective point-spread function (PSF) of the imaging, and estimates the level of spurious signals arising from e.g. detector imperfections. Our SOFIevaluator is model-free and unbiased in the sense that all of these parameters are determined purely from the acquired data, without the need for additional input.

In addition to presenting the fundamental algorithm, we also used SOFIevaluator to test the suitability of 20 different fluorescent proteins, spanning the full optical window, for SOFI imaging.

## 2. Metrics for a quantitative comparison of SOFI data

The SOFIevaluator calculates multiple parameters from a single acquired dataset, reflecting the difficulty in capturing the image quality of a (superresolved) fluorescence image in a single number. For example, signal-to-noise ratio (SNR) and spatial resolution are often presented as quality metrics for the imaging, but are incomplete in the sense that an image with high resolution can have a low SNR while a high SNR image might have a low spatial resolution.

The essence of our approach is the dependence between the fluorophore and detector properties, and the cross-cumulants calculated in SOFI imaging. This dependence is fully quantitative and well understood, reflected in the availability of both simplified and complete models of the imaging [2, 18]. We based our methodology on the calculation of second-order cumulants, corresponding to second order SOFI imaging, as these cumulants can be calculated quickly and represent a good balance between ease of calculation and information content.

A second-order cross-cumulant is calculated by considering the contributions of two different pixels on the detector (camera). In SOFI, a cross-cumulant gives rise to a virtual pixel that lies in between these two pixels [18]. A very large number of such cross-cumulants can be defined, reflecting different orientations and physical distances between the detector pixels. Similarly, cross-cumulants can also be constructed between detector pixels that are not only separated in space but also in time, meaning that the pixels values are taken from different images in the acquisition stack. These ‘time lags’ can take on only integral values: zero for pixel values from the same image, one for pixels from consecutive images, etc. By calculating cross-cumulants for different pixel combinations and different time lags, information about the spatial resolution as well as the temporal dynamics can be obtained from a single dataset.

Our SOFIevaluator makes full use of this additional information. In particular, the algorithm initially calculates second-order cumulants for increasing time lags to judge the contribution of bleaching to the signal, and to assess whether the camera acquisition speed matches well with the blinking dynamics of the label, as summarily introduced by Geissbuehler et al. [19]. Next, the algorithm estimates the signal-to-noise ratio using statistical resampling, based on our previous work [17]. Finally, using a theoretical formalism outlined in [18] and [2], SOFIevaluator estimates the point-spreading function (PSF) of the system by comparing the SOFI signal relative to the physical distance between the detector pixels. An overview of a typical SOFIevaluator output can be found in Figure 1. The source code for the SOFIevaluator is available at https://bitbucket.org/dedeckerlab/SOFIevaluator/.

**Fig. 1.**
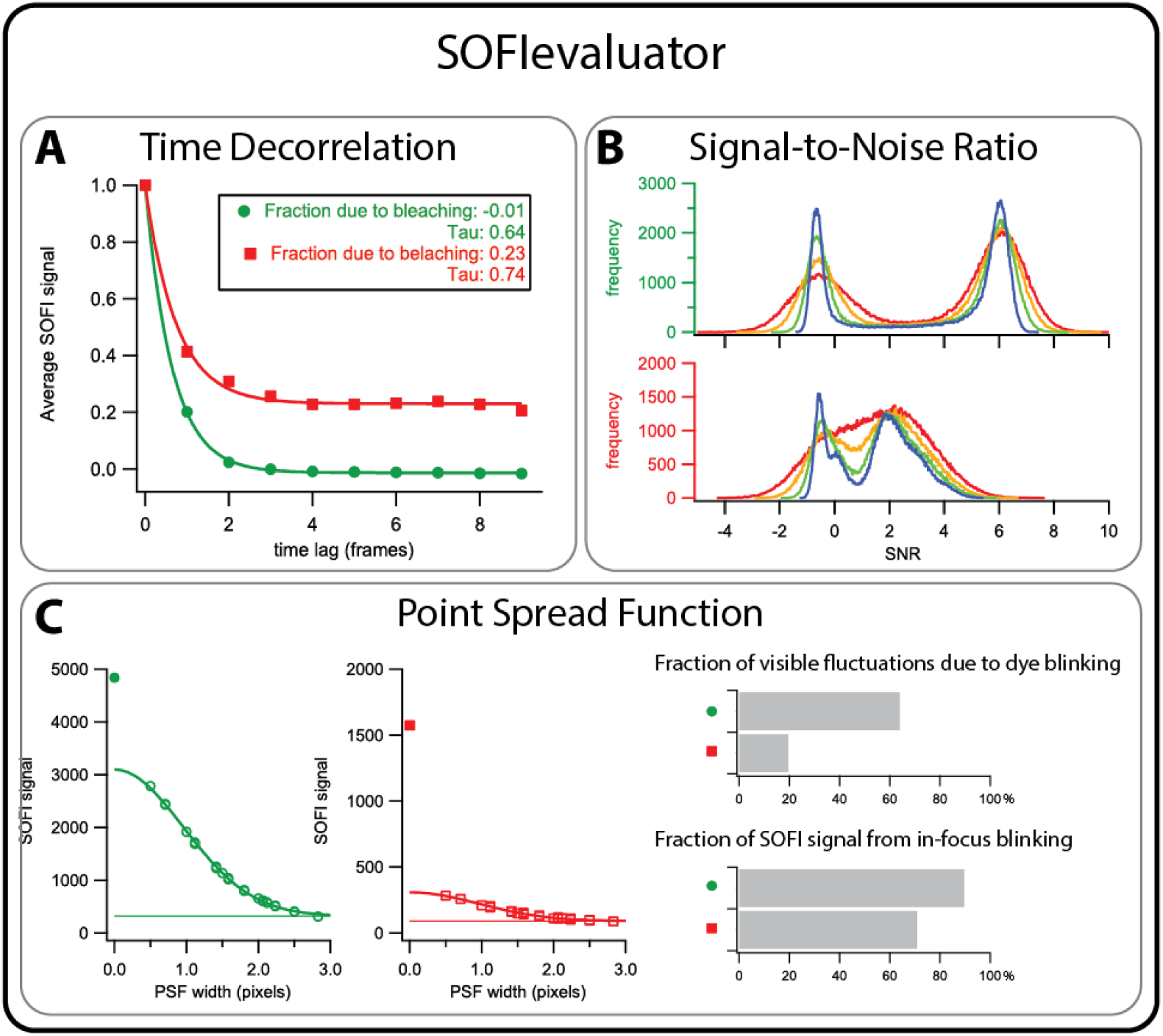
Overview of the SOFIevaluator output. Green spheres and the top graph in panel **B** represent data from SkylanS (imaging condition 2, images in Figure 2A). Red squares represent data from rsGreen1 (imaging condition 6). **C**: open markers: spatial decorrelations data points; filled markers: autocorrelation function; horizontal line: value of fit at infinite PSF width.

### 2.1. Determining blinking kinetics and the influence of photodestruction

One of the main determinants for SOFI quality is the rate of the fluorescence dynamics of the labels relative to the camera frame acquisition rate. While in theory it is possible to perform SOFI microscopy with a wide range of fluorophore blinking rates, in practice an unfavorable setting will reduce the quality of the images by slowing down the convergence of the SOFI image. In general, the average emissive duration (‘on-time’) of the fluorophores should roughly match the duration of a single camera exposure. This can be tuned by changing the acquisition rate of the camera, or alternatively by varying the power of the light irradiation for light-induced blinking processes.

In Ref. [19], an approach was introduced for estimating this blinking speed from the SOFI data. A so-called decorrelation plot is made of the average signal in a SOFI image against the time-lag used in the calculation. The time constant of an exponential fit to this curve is then considered to be the blinking period of the label. While our work has shown that this trend is not in general exponential in nature [2], it is often a very good approximation, and the resulting decay time (‘tau’) is a useful indicator of the blinking speed. Slow blinking translates to tau values higher than one frame, while fast blanking leads tau values below one. As a rule of thumb, optimal results can be expected when this decay time is close to one.

Photodestruction of the fluorophores has a pronounced effect on SOFI imaging, since it introduces correlations into the data and therefore leads to additional signal that does not contain superresolution information. As we have previously shown [20], its contribution results in images that appear to have a higher SNR, due to the increased signal derived from the photodestruction, yet that do not faithfully represent the underlying structure of the sample. In that work we also demonstrated that ‘batching’ of the images is an effective solution to this issue, meaning the calculation of multiple SOFI images from subsets of the data that are then added or averaged together.

In practice, however, it may not be clear what the influence of photodestruction, and hence the optimal batch size, is for a particular dataset. Our SOFIevaluator makes it possible to determine this. The key insight is that blinking is usually on a much faster time scale compared to photodestruction, and hence the cross-cumulants should rapidly decay to zero for even small lag times. The comparatively much slower bleaching will manifest as an additional long-lived contribution or offset in the decorrelation curve (Figure 1A). More generally, any process that causes pixels to be correlated over multiple images will introduce additional components into the correlation curve. Photodestruction is the most likely culprit, though effects such as instability of the excitation source will also show up here. The batch size used in the calculation can then be adapted to minimize its effect.

### 2.2. Quantifying the SNR of a SOFI image

We previously introduced a general framework for determining the uncertainty on any pixel in a SOFI image using statistical resampling [17]. Using this formalism, we can readily calculate the SNR of any single pixel by dividing its value by the associated standard deviation.

In practice, however, such per-pixel SNR estimates are highly variable because most images are highly heterogeneous, with some parts of the image showing high fluorescence brightness, others moderate brightness, and yet others containing only background. Reducing this heterogeneity into a single metric for the SNR, that can then be compared across datasets and acquisitions, is not a simple task.

Our standard approach is to calculate all of the per-pixel SNRs and to visualize these in a histogram. The resulting histogram then consists of a peak centered at zero, showing background pixels, and one or more additional peaks showing the SNR associated with image features (Fig. 1**B**). The image SNR is then reported as the maximum of this peak or peaks.

In practice, the background peak is sometimes present at a value somewhat below zero, resulting from a very small negative offset in the signal of background pixels that arises in the calculation. The ratio of this offset with the very small noise that is associated with these pixels, leads to an SNR value that becomes negative. An SNR histogram with a single peak at or below zero is indicative of very little, if any SOFI signal, with most pixels being background.

SOFI datasets containing relatively few fluorescence images, or noisy images, may cause the SNR histogram to broaden. In this case one can apply smoothing of the calculated per-pixel SNR data, which reduces the noise yet leaves the peak maxima largely unchanged (Fig. 1**B**). The SNR can then likewise be determined as the maximum of the signal peak.

### 2.3. Estimating the effective PSF of SOFI images

In SOFI, one tries to calculate second-order cross-cumulants between two detector pixels that each show intensity variations due to fluorophore blinking. Since this blinking is assumed to be independent, non-zero cross-cumulants for blinking processes can arise only when the same fluorphore emits in both detector pixels simultaneously, which is determined by the size of the PSF. The calculated cross-cumulants become smaller and smaller in magnitude as the distance between the detector pixels increases, since it becomes increasingly unlikely that a single fluorophore will be detected by two pixels that are further and further apart. Ultimately, if the SOFI signal arises purely through blinking, the calculated cross-cumulants will be zero for pixels that are further apart than the extent of the PSF.

The resulting distance dependence of the SOFI signal can be readily calculated for any experimental dataset, simply by selecting appropriate cross-cumulant combinations (Figure 1**C**), yet provides direct information on the quality of the measurement. First of all, the width of this curve is a factor 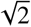 larger than the width of the native instrument PSF [18]. Rescaling the curve by this factor, as we did in Figure 1**C**, should therefore deliver the classical widefield PSF. If these do not agree sufficiently well (given noise) with the expected value determined using e.g. fluorescent beads, there are likely issues with the quality of the input data, and the resulting SOFI image should not be trusted fully. The actual SOFI second order PSF is an additional factor of 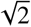 more narrow.

In actual measurements, the spatial decorrelation graph may not fall off to zero for large pixel separations (Figure 1**C**). This means that there is also SOFI signal present from origins other than in-focus emitter blinking. Two further benign components are usually to blame. Firstly, since all blinking molecules contribute to the PSF estimation, out-of-focus emitters will lead to a broadening of this curve. Secondly, emitter bleaching also manifests itself as long-distance correlations, though the contribution of this is best verified using the temporal decorrelation strategy presented above, and corrected by batching the images in the calculation [20]. However, long-distance contributions may arise also from artificial correlations between pixels, due to the imperfections in camera design or construction. In practice, these effects are small for well-corrected scientific cameras [16].

Finally, we note that this analysis allows direct estimation of the contribution of noise to the visual dynamics observed in the sample. When visually inspecting a set of acquired fluorescence images, it can be difficult to visually distinguish emitter blinking from the apparent emission dynamics arising from shot noise or other noise sources. Since this noise does not correlate between different pixels, their contribution results in an increased signal when the SOFI cumulants are calculated for a pixel distance of zero, that is, within the same detector pixel. The fraction of visible fluctuations in the data that is due to bye blinking can then be determined by comparing the increased amplitude of this autocumulant with that extrapolated from the (unaffected) cross-cumulants (Figure 1**C**).

## 3. Evaluating the suitability of selected fluorescent proteins for SOFI

We next sought to apply our evaluation framework to determine the suitability of selected fluorescent proteins for SOFI imaging. To achieve this, we fused the membrane targeting domain of Lyn kinase N-terminally to each label, directing it towards sphingolipid- and cholesterol-enriched microdomains of the plasma membrane [21] in Cos7 cells. In the blue region, EBFP2 [22] and mTagBFP2 [23] were used while cyan proteins were mCerulean [24], mTurquoise2 [25], and mTFP0.7 [26]. In the green wavelength region, wQ [27], rsGreen1 [28], EGFP, Dendra2 [29], SkylanS [30], ffDronpa [31], mNeonGreen [32], mEos3.2 [33], and EYFP were used. Dendra2 and mEos3.2 were chosen because they are also excellent probes for photoconversion-based PALM. In the orange to red region, we chose mOrange2 [34], mKO2 [35], rsTagRFP [36], mScarletI [37], rsFusionRed3 [38] and mCherry. Some photophysical properties of these probes are given in Table 1.

**Table 1.** Labels analyzed with SOFIevaluator. λ_Ex_ = excitation maximum (nm), λ_Ex_ = emission maximum (nm), B = molecular brightness, PT = phototransformation properties, PS = photoswitchable, PC = photoconvertable

| Label | $\lambda_{Ex}$ (nm) | $\lambda_{Em}$ (nm) | B | PT | laser |
| --- | --- | --- | --- | --- | --- |
| EBFP2 | 383 | 448 | 17.9 |  | 405 |
| mTagBFP | 399 | 454 | 32.8 |  | 405 |
| mCerulean | 433 | 475 | 16.2 |  | 445 |
| mTurquoise2 | 434 | 474 | 27.9 |  | 445 |
| mTFP0.7 | 453 | 488 | 30.0 | PS | 445 |
| wQ | 476 | 504 | 25.8 | PS | 488 |
| rsGreen1 | 487 | 511 | 25.6 | PS | 488 |
| EGFP | 488 | 507 | 33.5 |  | 488 |
| Dendra2 (green) | 490 | 507 | 22.5 | PC | 488 |
| SkylanS | 499 | 511 | 97.5 | PS | 488 |
| ffDronpa | 503 | 518 | 78.8 | PS | 488 |
| mNeonGreen | 506 | 517 | 92.8 |  | 488 |
| mEos3.2 (green) | 507 | 516 | 53.3 | PC | 488 |
| EYFP | 513 | 527 | 44.9 |  | 488 |
| mOrange2 | 549 | 565 | 34.8 |  | 561 |
| mKO2 | 551 | 565 | 39.6 |  | 561 |
| rsTagRFP | 567 | 585 | 4.1 | PS | 561 |
| mScarletI | 569 | 593 | 56.2 |  | 561 |
| rsFusionRed3 | 580 | 607 | 3.0 | PS | 561 |
| mCherry | 587 | 610 | 15.8 |  | 561 |

Each of these labels was imaged under 12 different imaging conditions (see Materials and Methods and Figure 3). Briefly, we acquired image stacks with varying levels of excitation intensities, accompanied with different intensities of 405-nm light. For each condition, 10 different fields of view approximately the size of one cell were imaged. The resulting 2240 image stacks of 500 images were then analyzed using the SOFIevaluator.

To analyze the resultant quality assessment parameters, we removed individual datapoints that had a modified Z-score of above 3, which means they can be considered outliers [39]. The origin of these outliers might be very diverse: biological (e.g. cells express at different levels and sometimes show aberrant phenotypes), instrumental (e.g. cell that are slightly out of focus, more than one cell in a given field of view, very bright background speckles…) or due to the analysis software (e.g. fits that did not correctly converge).

We settled on the SNR peak as the main quality criterion, as we found it to be robust and best describing the visual quality of the images. Secondarily, we judged the other parameters, giving extra insight into which imaging conditions might be best for a given label. Summarizing graphs of the SOFIevaluator output are given as Supplementary Data 1.

### 3.1. Blue labels

We tested two blue fluorescent proteins, EBFP and mTagBFP, neither of which is reported to show switching or blinking behavior. The PSF FWHM was around 2.4 px (EBFP2) and 2.1 px (mTagBFP). This is slightly larger than the theoretical value of 1.9 px, which we can attribute to instrumental imperfections. Although we see that increasing the excitation light increases the SNR of the image, the average SNR maxed at 0.24 (EBFP2) and 0.47 (mTagBFP). On average less than 33% (EBFP2) and 13% (mTagBFP) of the signal being due to bleaching, so the low SNR is certainly not derived from an excessive bleaching contribution. However, >90% of the visible fluctuations in the image is not due to dye blinking, and from the remaining <10%, only half is derived from in-focus blinking. In other words, the blue FPs display very low levels of SOFI-compatible blinking dynamics under the conditions tested here. It is thus safe to say that none of the blue fluorescent proteins that we tested is suitable for pcSOFI, at least not under the conditions that we tested.

### 3.2. Cyan labels

We tested 3 cyan fluorescent proteins: mCerulean, mTurquoise2 and mTFP0.7. All three showed a PSF FWHM of 2.1 to 3.0 px which is acceptable, given the theoretical value of 2.0 px at 480 nm emission. However, mTFP0.7 can immediately be discarded as a suitable pcSOFI label under our imaging conditions, since the SNR is consistently below 0, indicating that most of the apparent signal is indiscernible from the imaging background. For mCerulean and mTurquoise2, the SOFI data contains a very high contribution of bleaching (40% and higher) and a very high fraction of the signal (>80%) are fluctuations that are not due to dye blinking. From the fluctuations that do contribute, less than half of the fluctuations results from in-focus blinking molecules. This means that mCerulean and mTurquoise2 provide, under our experimental conditions, very little actual blinking. The high fraction of the signal that is due to bleaching suggests using a smaller batch size for the SOFI calculation, though we found that re-evaluating the data with smaller batch size (25 instead of 50) did not result in improved quality (data not shown). As a result, we could not identify a suitable cyan fluorescent protein for pcSOFI imaging.

### 3.3. Green labels

We tested nine green fluorescent proteins for fitness with pcSOFI imaging (Table 1). After removal of the outliers, almost all PSF FWHMs were located between 2.1 and 2.5 px, which is in good agreement with the expected value at these emission wavelengths (2.1 px).

We find that most most labels, with the exception of wQ, perform well in terms of SNR, with SNR peaks of on average between 3 and 6. Especially the GFP-derived FPs (EGFP, rsGreen1 and wQ) benefit from a high illumination intensity. In this set, SkylanS is most noteworthy, as the spread in position of the SNR peak across the 10 imaged cells is reasonably small (except for the high 405 nm illumination intensity conditions). This is indicative of minimal biological and instrumental variability that, together with good compatibility with SOFI imaging, leads to robust data and analysis. With an SNR distribution peak of on average 5.8 and almost no contribution of bleaching to the SOFI signal, condition 2 (see Figure 3 for the imaging conditions) of SkylanS is performing remarkably well.

A representative image of this imaging condition, as well as images of the best- and worst-SNR SkylanS-expressing cells imaged with condition 8 can be found in Figure 2. As can be seen, biological and instrumental variability can have a large influence on the acquired parameters and the results obtained with the SOFIelvaluator are not a criterion for the subjective aesthetic quality of the images.

**Fig. 2.**
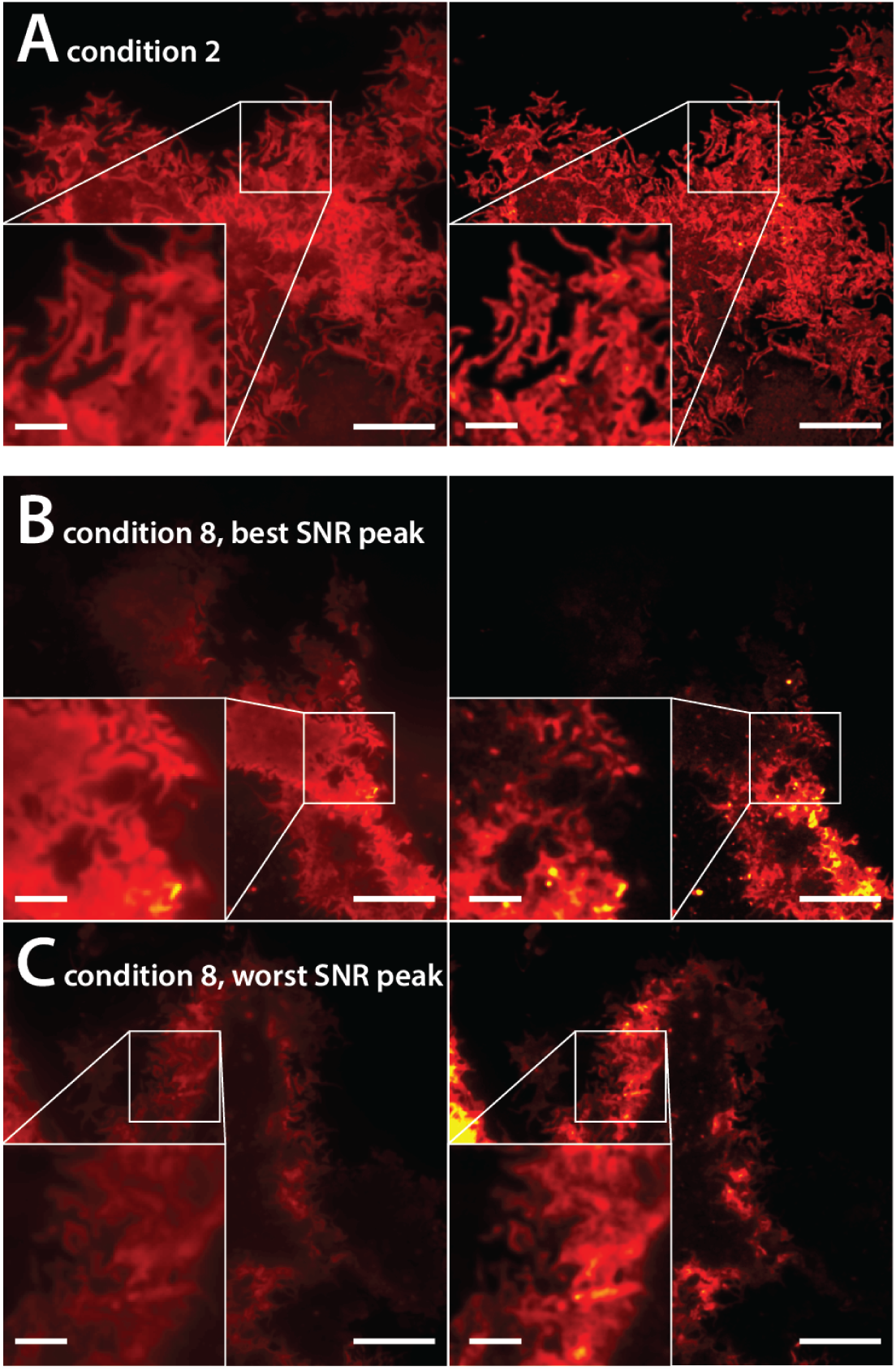
Average and SOFI images of lyn-SkylanS-expressing cells. (A) Representative cell imaged under imaging condition 2. (B-C) Cells with the best (B) and worst (C) SNR peak of the 10 cell imaged under imaging condition 8. Left = average image, right = SOFI image. Scale bar = 10 µm (large image) and 2.5 µm (inset)

Note that, under illumination condition 2, similar quality parameters can be obtained for mEos3.2 (the parent of SkylanS) and ffDronpa as for SkylanS. The SOFI images obtained with these labels are equally good under the conditions that we tested here. An advantage of these labels is that they show photoswitching behavior (which is moderate, though present in mEos and its derived proteins [40]), the dynamics of which can be tuned by modifying 488- or 405-nm light intensities, thus modifying the blinking behavior to match the SOFI algorithm best, yielding high quality SOFI images.

### 3.4. Red labels

All red labels that we tested seem to reach a PSF FWHM of around 2.5 px, which is close to the expected value of 2.4 px. We see a big spread in the location of the peak of the SNR distribution. For all but mScarletI, the peak is located close to or below zero for the lower illumination intensities. This is indicative of a lack of SOFI signals and prevalence of pixels that are background or indiscernible thereof.

mScarletI and mCherry perform on par in terms of the fraction of signal that is a result of photobleaching (+-60%) and the fraction unexpected SOFI signal (+-20%). Redoing the analysis with smaller batch sizes might reduce this unwanted bleaching contribution. Again, as with the green labels, higher illumination intensity is correlated with better quality parameters. Overall, the red labels perform better than the blue or cyan-emitting FPs, but are outperformed by the better green labels.

Interestingly, we previously found that rsTagRFP yielded informative images [41], while the SOFIevaluator suggests poor SOFI image quality using this label. Using the equipment used here, the lower molecular brightness of rsTagRFP (and rsFusionRed3) results in images that are purely noise with barely-visible cells, especially for the low excitation intensities that we tested. This is well reflected in the SNR peak that is at or below zero, but increases as excitation and 405-nm intensities increase. Consequently, most of the visibly observed intensity fluctuations (>90%) are not a result of photoblinking. Together with the fairly low contribution of photobleaching to the SOFI data (<5%), this suggests that rsTagRFP and rsFusionRed3 might perform better at higher illumination intensities. We re-analyzed our 2012 data on rsTagRFP ([41], Fig. 6) using SOFIevaluator and found parameters indicative of good SOFI data (SNR peak at 2.3).

We conclude that, under the imaging conditions that we tested, mScarletI functions well. Additionally, rsTagRFP and possibly rsFusionRed can be used as good labels for SOFI, though with higher excitation intensities than the conditions used here.

## 4. Conclusion

In this work, we developed a rigorous mathematical framework for estimating the quality of SOFI-generated images using 3 different paradigms: model-free SNR estimation, time decorrelation to estimate the contribution of bleaching to the signal, and estimation of the observed PSF. These tools represent a range of metrics that can be used to determine the quality of SOFI images. We suggest using the observed PSF width to determine whether the data are suitable for SOFI, followed by inspection of the obtained SNR to judge the overall image quality. The resulting software implementation is freely available at https://bitbucket.org/dedeckerlab/SOFIevaluator/.

To demonstrate the power of this evaluation tool, we used it to determine the suitability 20 selected fluorescent proteins for SOFI imaging, spanning much of the visual range. Using SNR as the main quality criterion, we concluded that the blue and cyan labels that we tested display poor compatibility with SOFI imaging, at least not under the conditions that we tested. We found that the red- (mScarletI and mCherry) and especially in the green-emitting FPs (SkylanS, ffDronpa and mEos3.2) function robustly well, especially with moderately low 488 nm light (green labels) and high 561 nm excitation light (red ones). Addition of 405 nm light in general did not lead to increased quality of the SOFI images.

In conclusion, we have developed an analysis strategy and framework that allow the unbiased estimation of the quality and reliability of a particular dataset for SOFI. By virtue of being model-free, our method can be applied to any acquired dataset, and does not require prior knowledge or familiarity with the analysis process. By introducing robust quality metrics into SOFI imaging, our work directly contributes to the development of robust and quality-assured super-resolution imaging in the life and materials sciences.

## 5. Materials and Methods

### 5.1. Calculations

The theoretical SOFIevaluator model was implemented in Igor Pro 8.03 (Wavemetrics). The code can be found on https://bitbucket.org/dedeckerlab/SOFIevaluator/.

### 5.2. Cloning and sample preparation

Plasmids were created by ligating PCR-amplified coding sequences for the different fluorescent proteins into a BamHI- and EcoRI-digested pcDNA3-lyn construct as described before. [28] All constructs were sequence-verified.

For imaging, 300 000 Cos7 cells were seeded in complete medium into a 35mm glass-bottom dish (P35G-1.5-14-C, MatTek). The next day, each dish was transfected with 1.5 µg plasmid DNA and 3 µL of FuGENE HD (Promega) following the manufacturer’s protocol. The day after, cells were washed once with and imaged in HBSS.

### 5.3. Imaging

The setup consisted of a Nikon Ti2 body equipped with a Nikon 100× CFI apo TIRF objective (NA = 1.49). Excitation light was provided by Oxxius 405, 445, 488 and 561 nm lasers through an Oxxius wavelength combiner box. The microscope is equipped with a ZT405/488/561/640rpcv2 dichroic and ZET405/488/561/640m emission filter for blue, green and red labels and a 440/488/561/635rpc dichroic with ET480/40m emission filter for cyan labels (all dichroics and filters from Chroma). The optical fibre was connected to the microscope through a manual Nikon TIRF illumination module and all image aquisitions were done in TIRF. Laser intensities were set at 3, 10, 30 and 100%, which resulted in 0.22, 0.96, 3.2 and 11.7 mW (405 nm), 0.3, 1.2, 5.0 and 14.1 mW (445 nm), 1.1, 4.1, 12.9 and 48.3 mW (488 nm) and 0.52, 1.7, 4.8 and 14.2 mW (561 nm); all intensities measured in epi mode. For additional 405 nm light stimulation in the cyan, green and red image channel, the 405 nm laser was set at 0, 1 or 10%, resulting in 0.000, 0.00115 and 1.115mW (cyan) and 0.000, 0.010 and 1.1mW (green and red). These conditions are labeled from 0-11 and summarized in Figure 3. Images were acquired with an Andor iXon 897 EMCCD camera operating at 33 Hz and an EM gain of 60 at 60 ^◦^C. The pixel size was 107 nm.

**Fig. 3.**
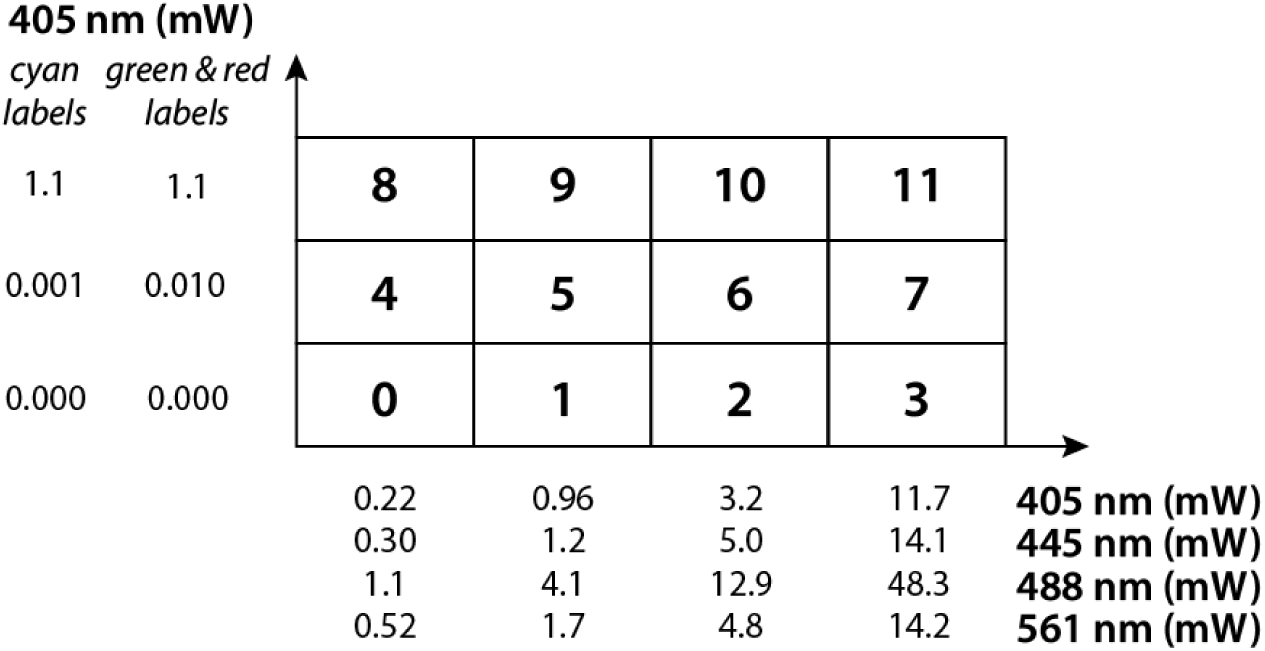
Summary of the imaging conditions

We set up an image sequence, whereby for each cell 4x 500 images were acquired, with increasing intensity of excitation light. This was repeated for each intensity of 405-nm light irradiation, leading to 30 cells imaged for each label. Exceptions are the blue labels, were only 10 cells were imaged (since the 405-nm light is the excitation light). Care was taken to select cells with brightness that was neither extremely high nor extremely low and preference was given to cells that were not in contact with other cells.

## Acknowledgments

The authors thank Sam Duwé for inspiring and fruitful discussions. B.M. and S.D. hold a postdoctoral fellowship from the Research Foundation-Flanders (FWO Vlaanderen). P.D. thanks the FWO Vlaanderen for funding via grants G0B8817N and G090819N and the ERC for funding via ERC Starting Grant 714688 (NanoCellActivity).

---

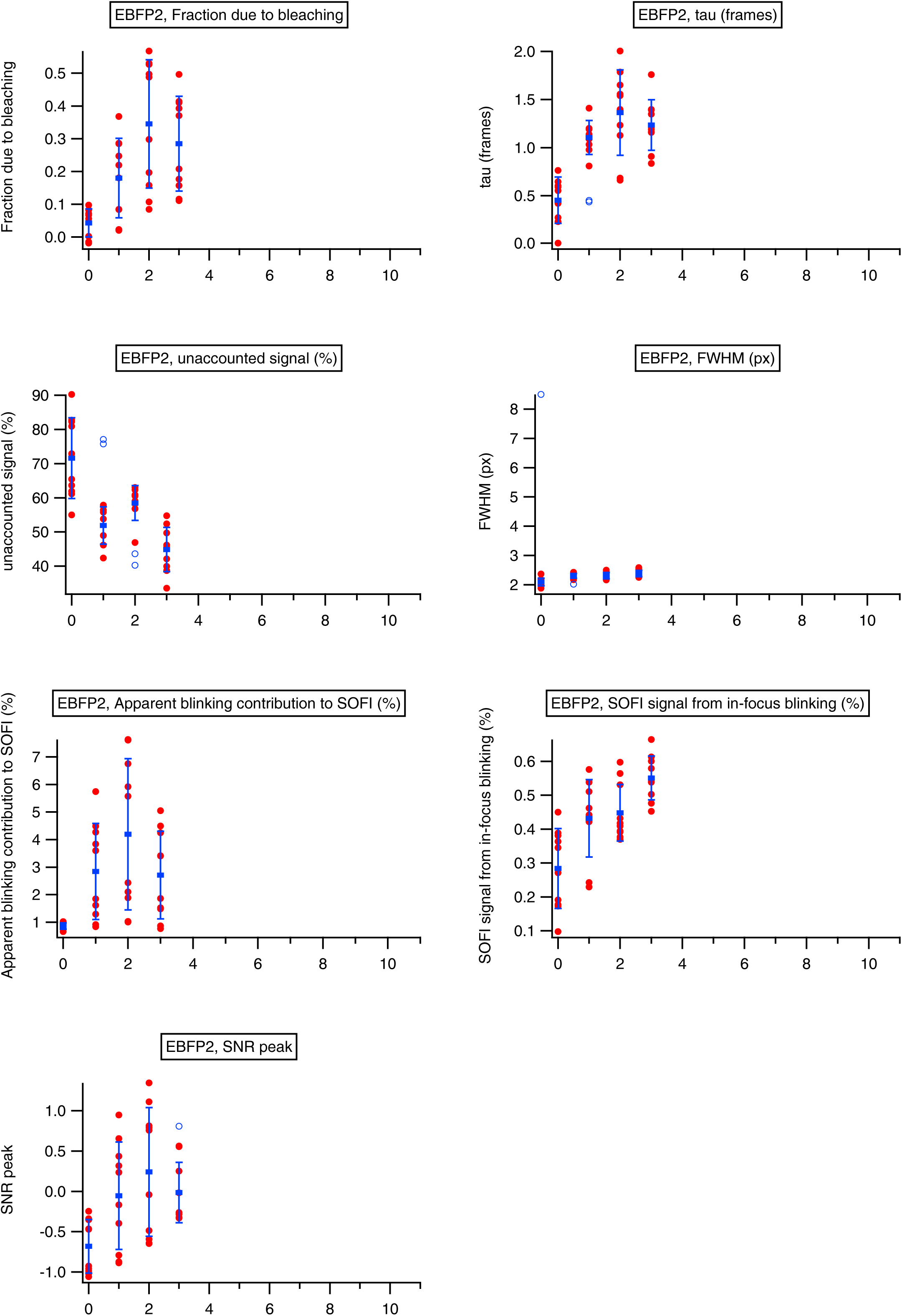
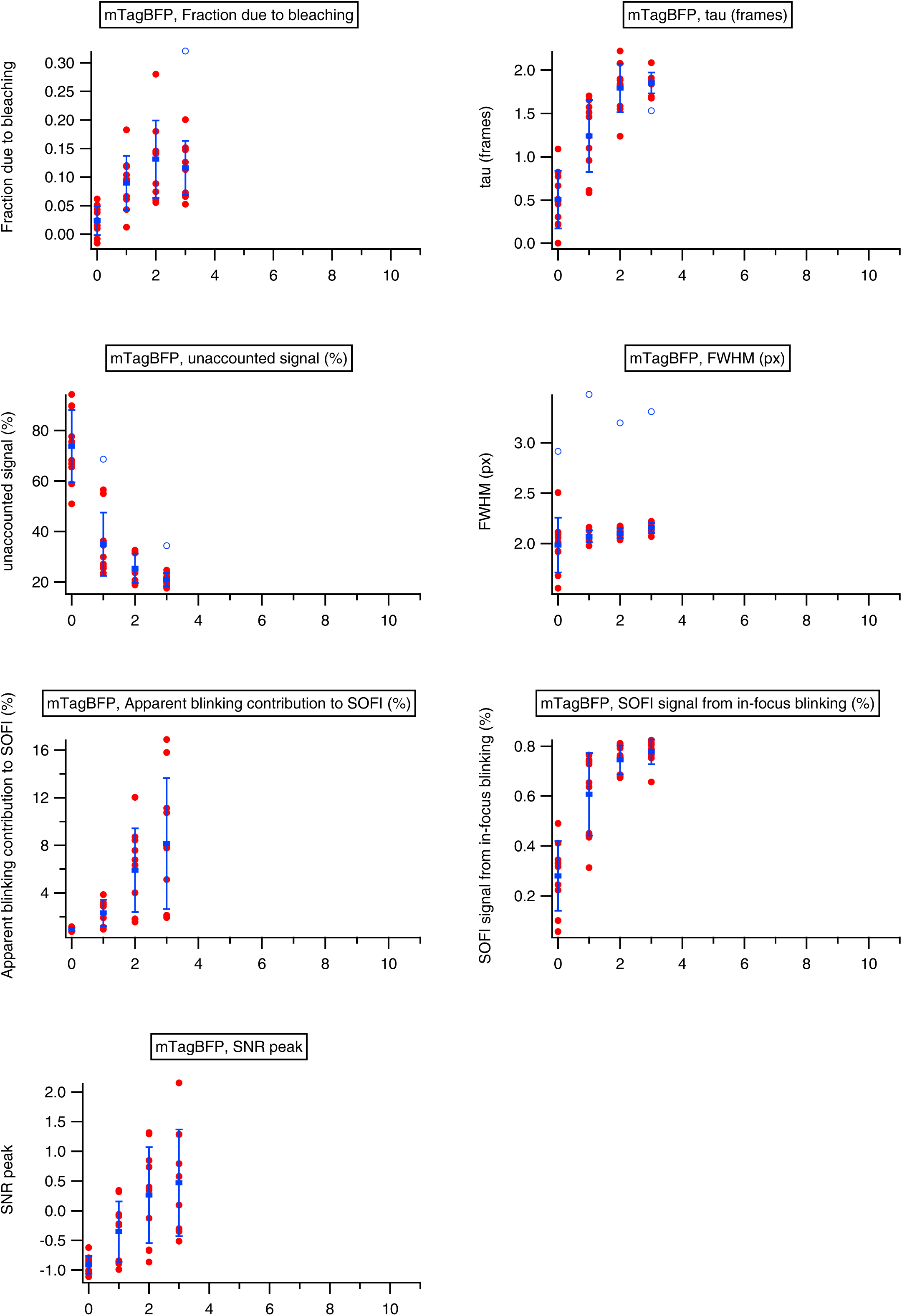
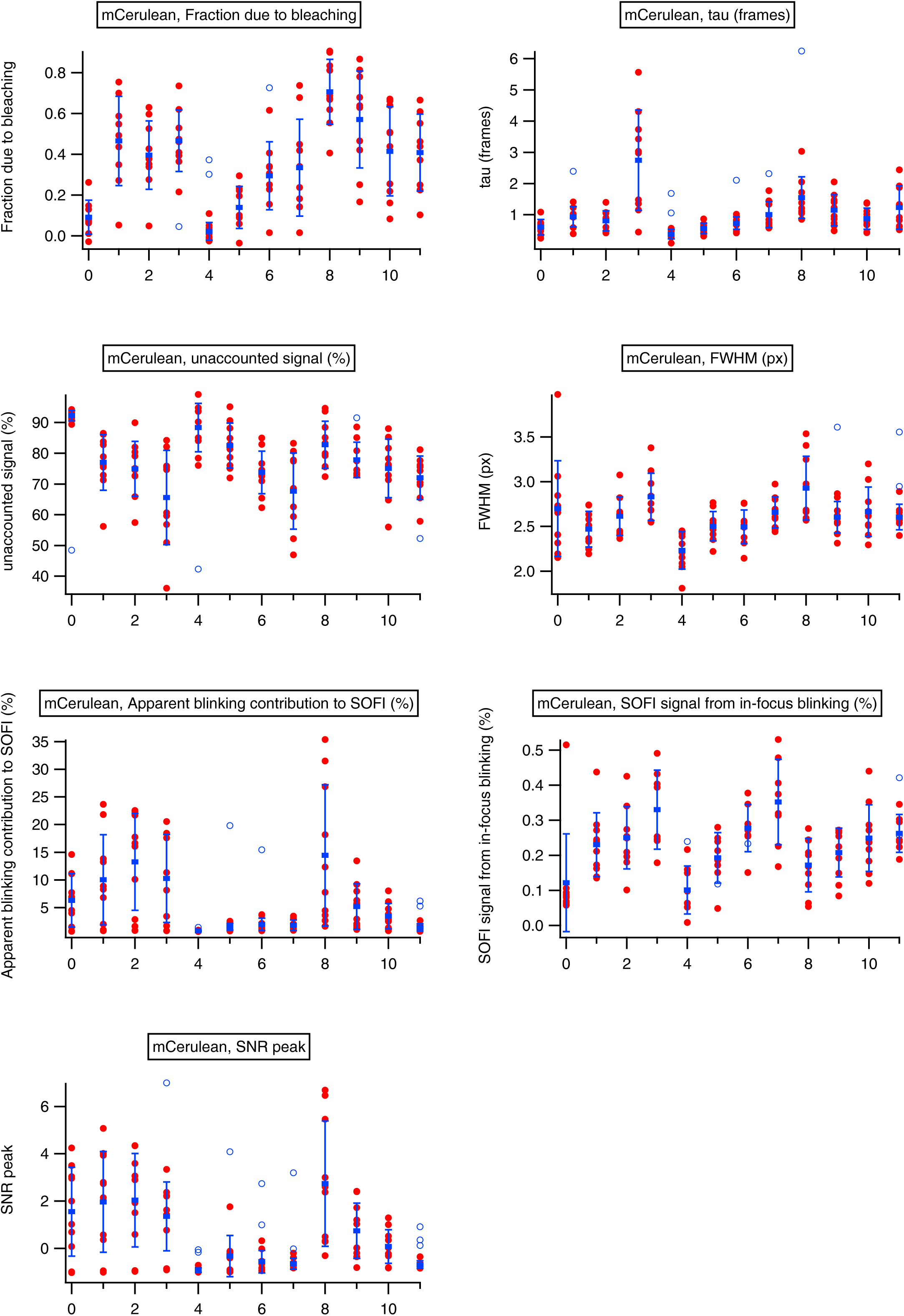
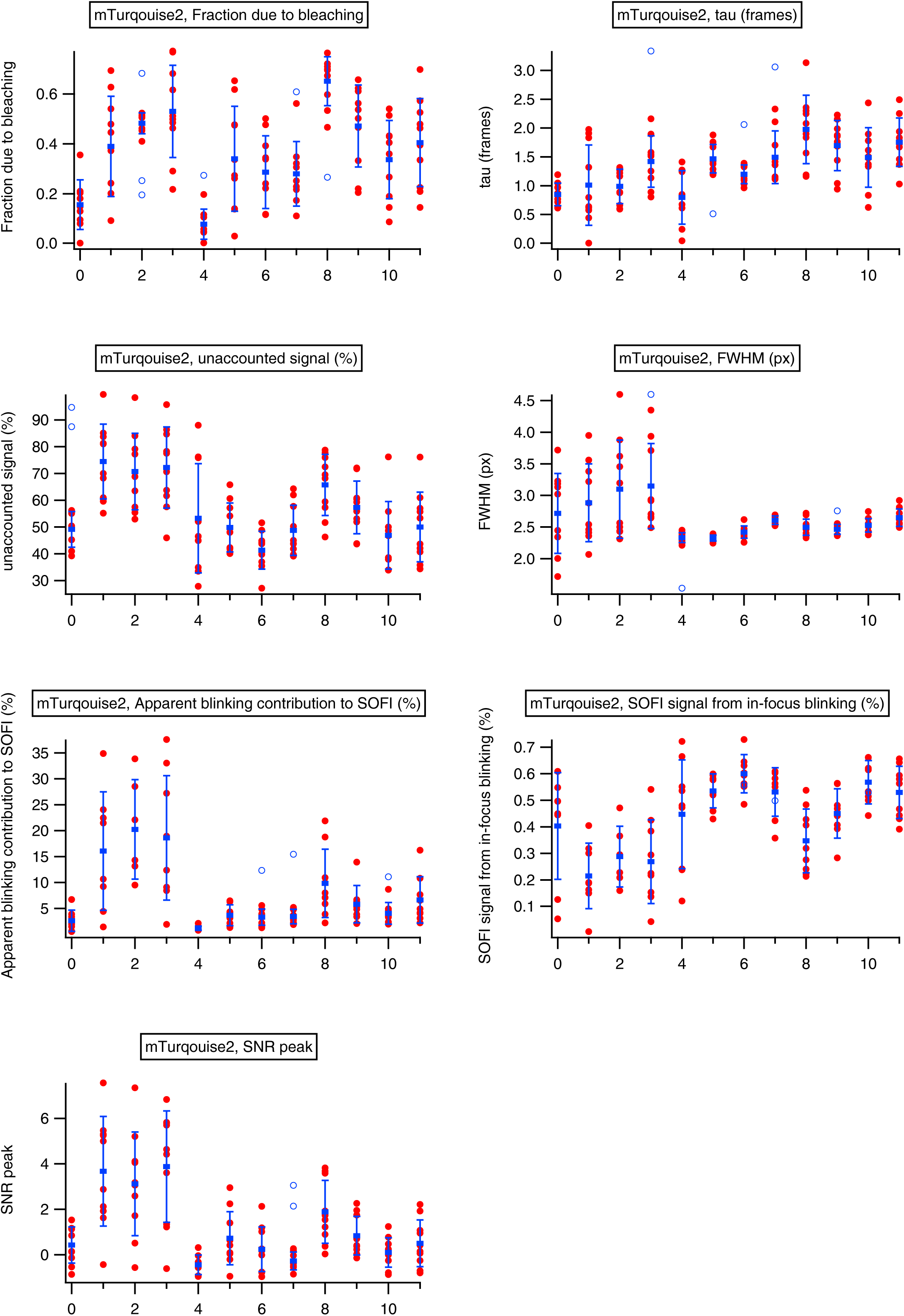
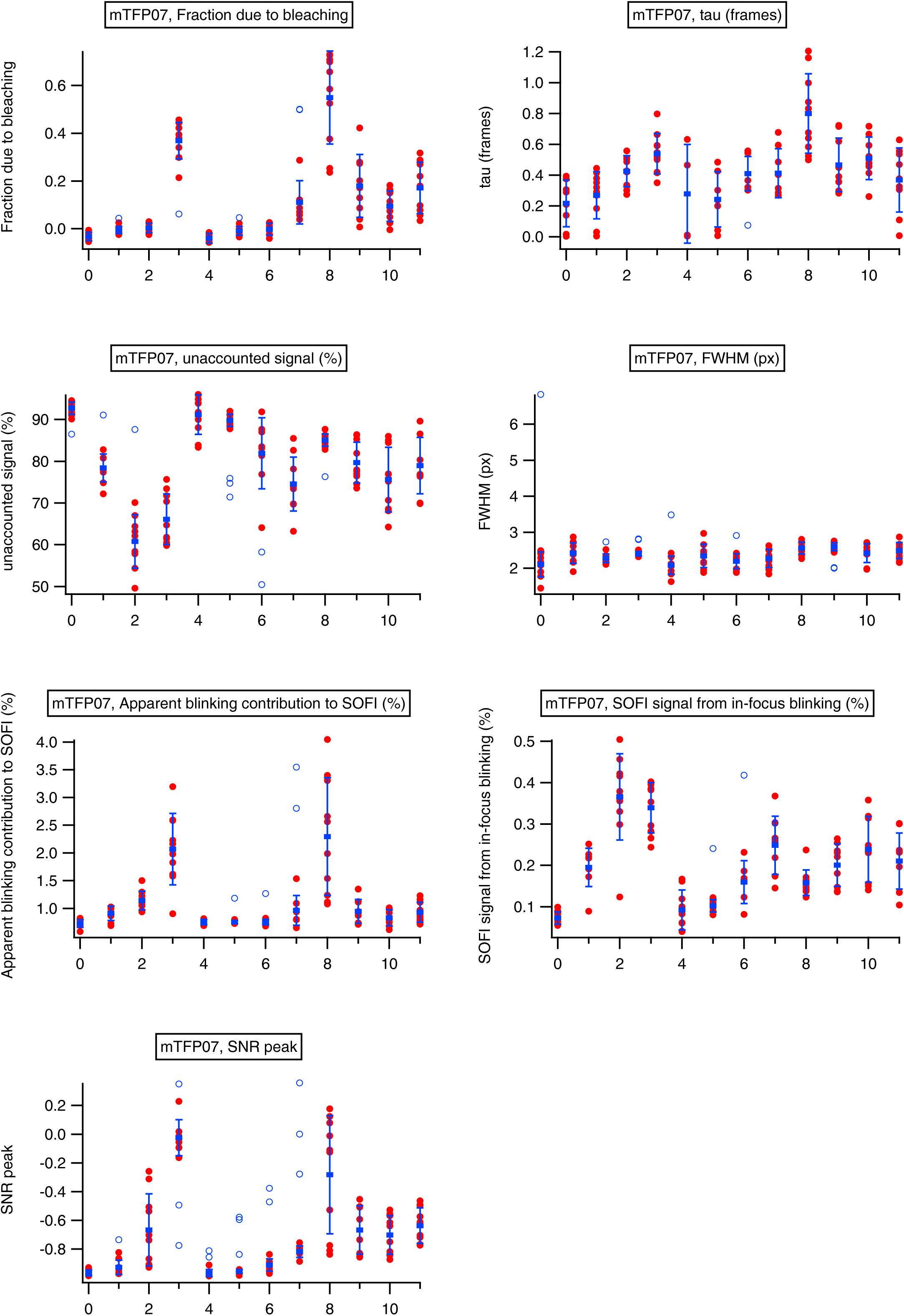
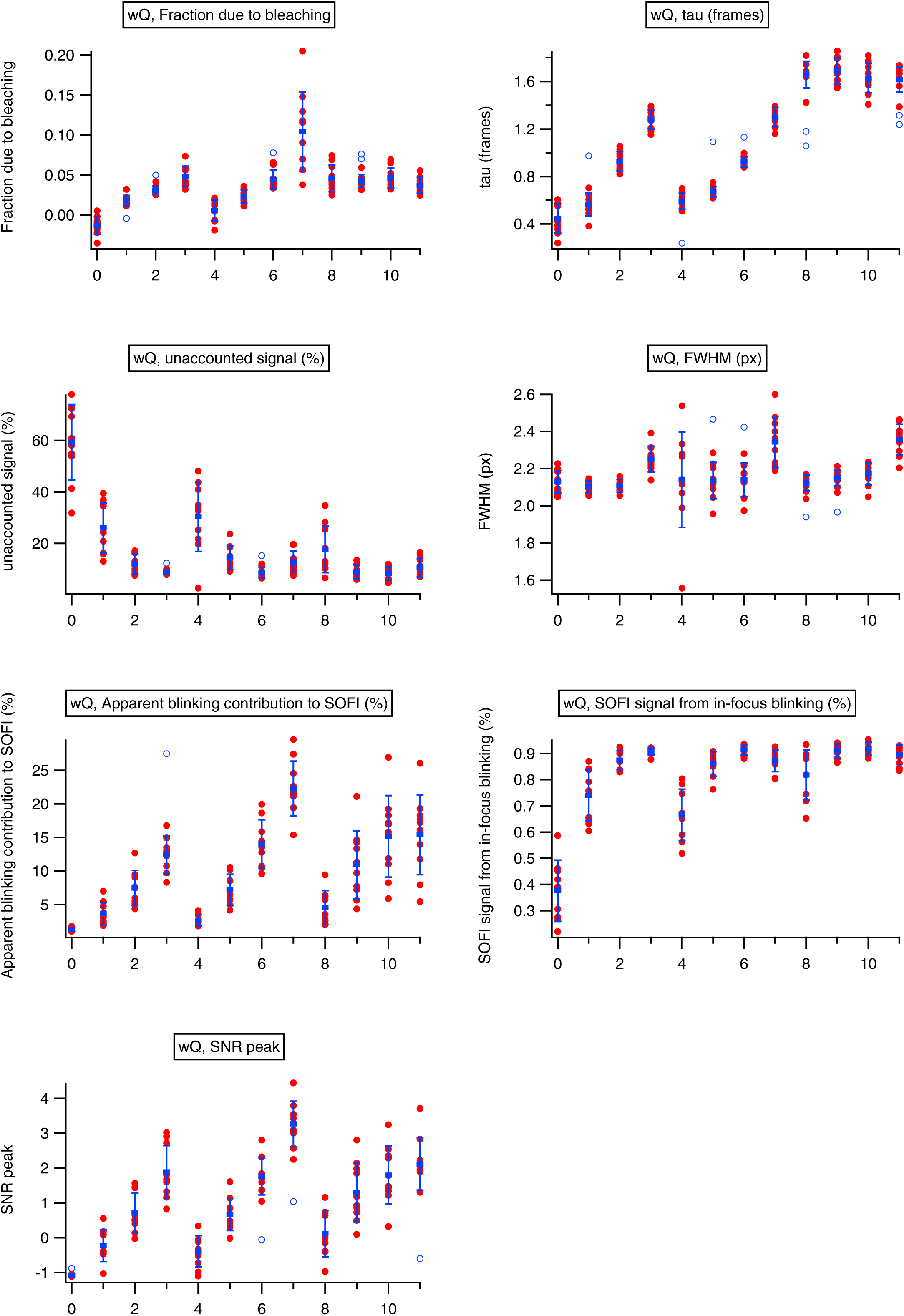
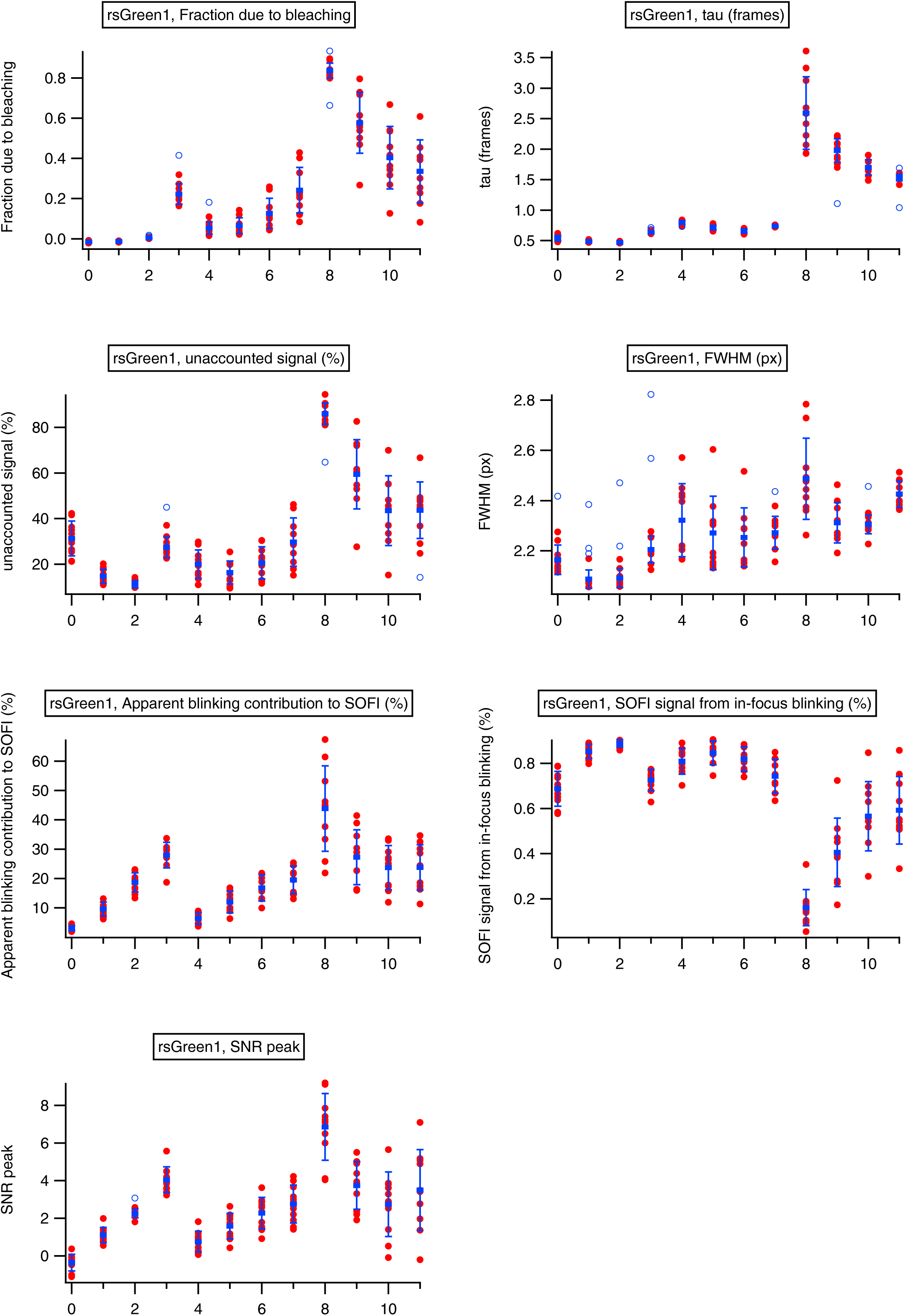
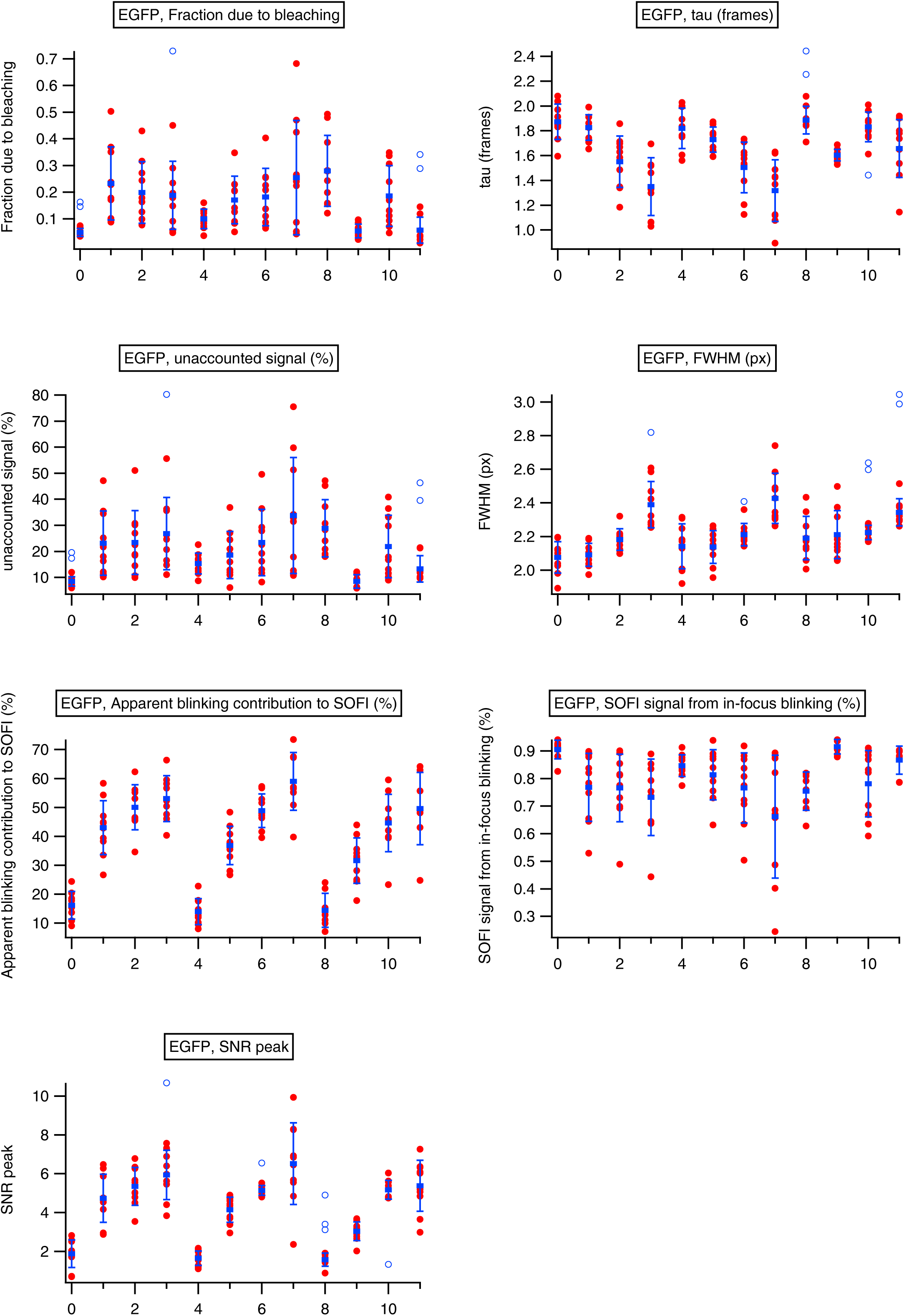
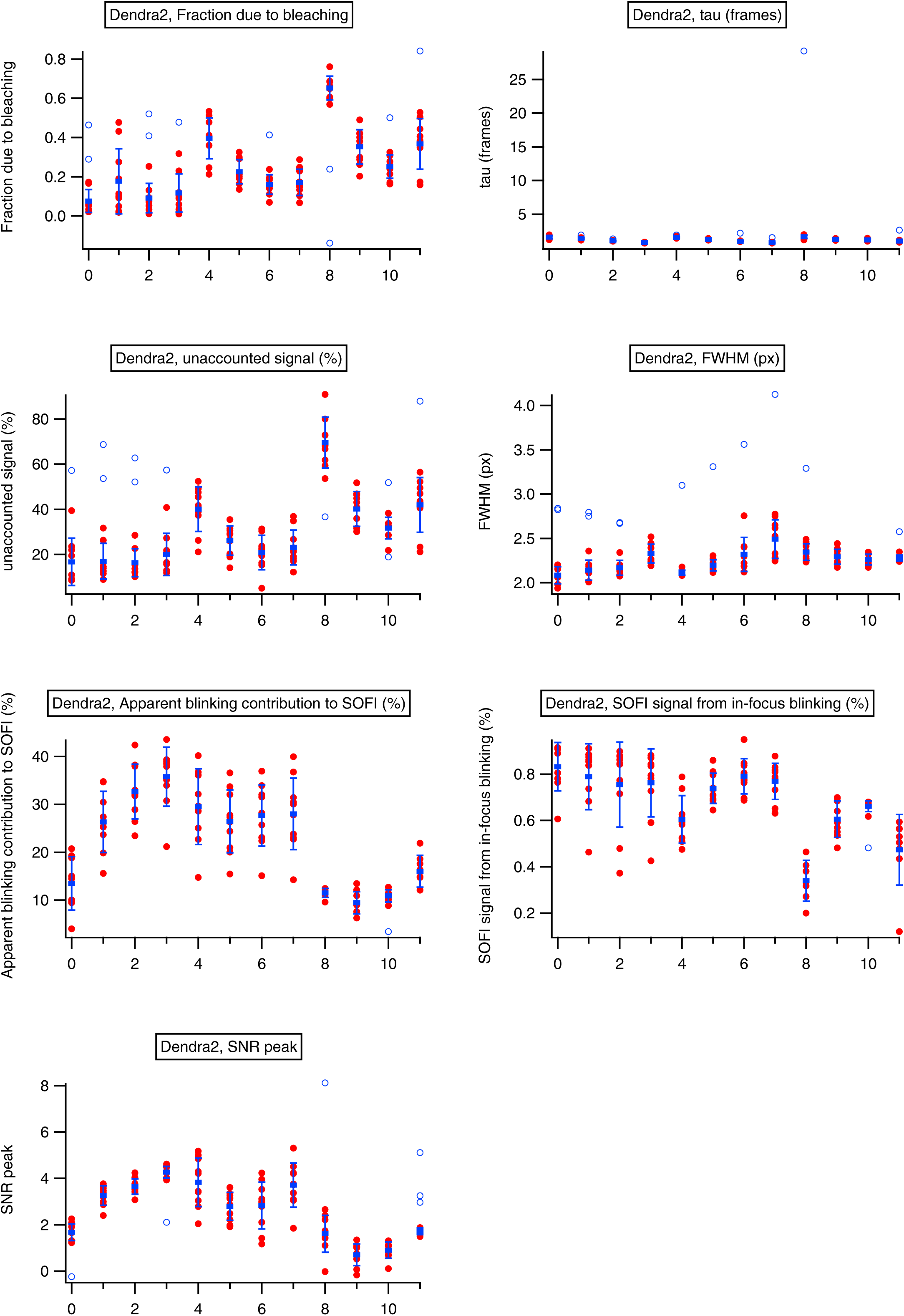
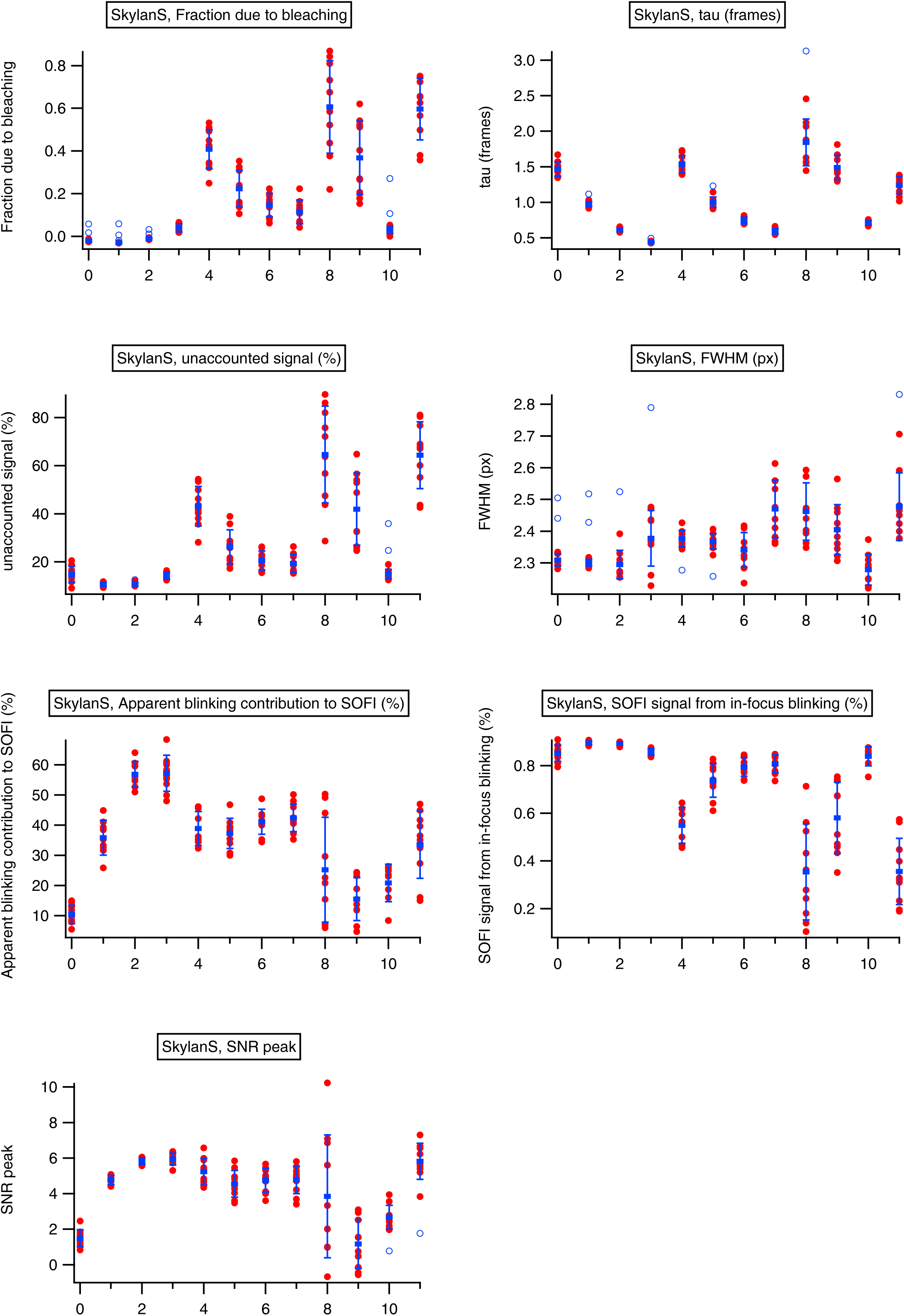
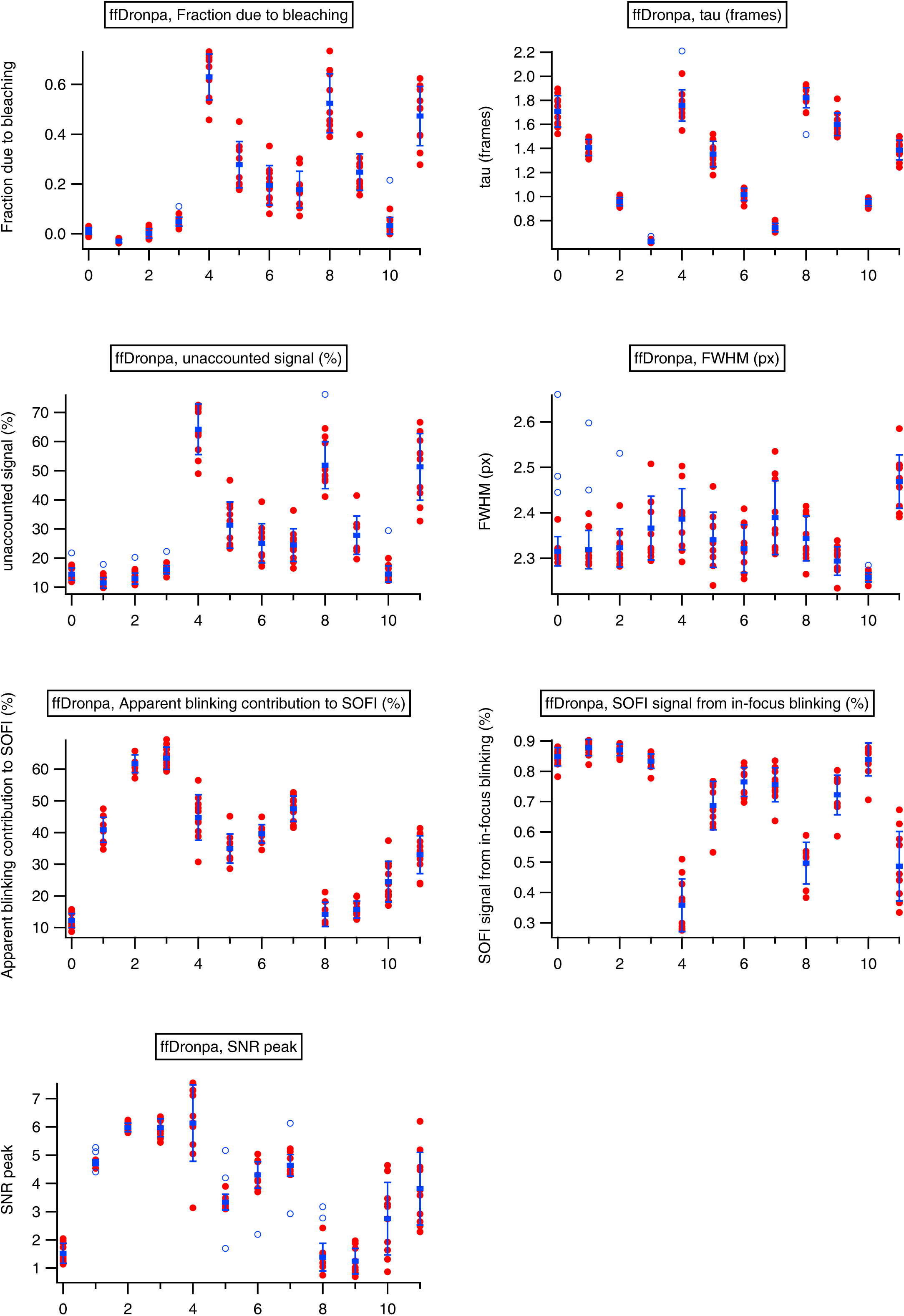
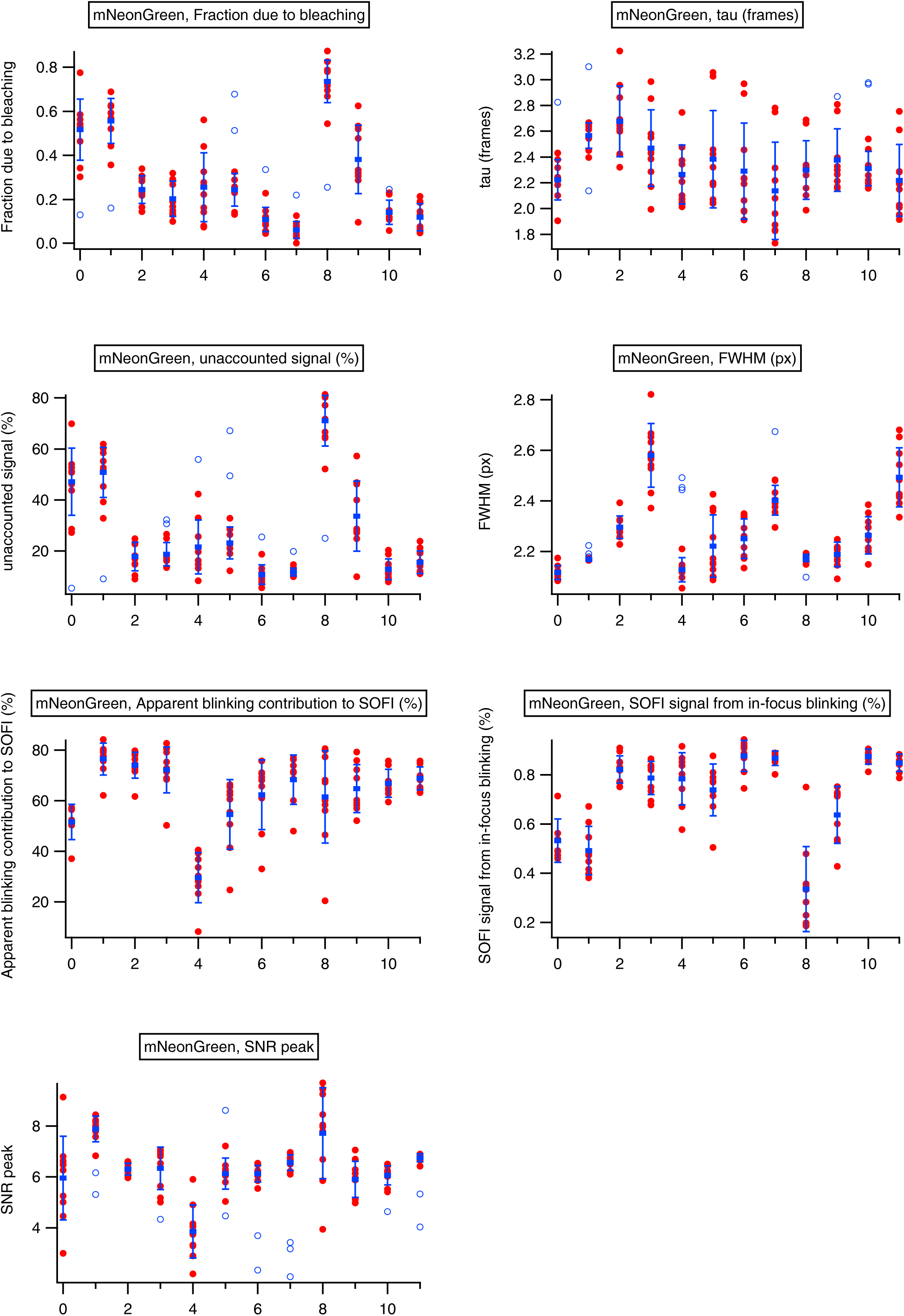
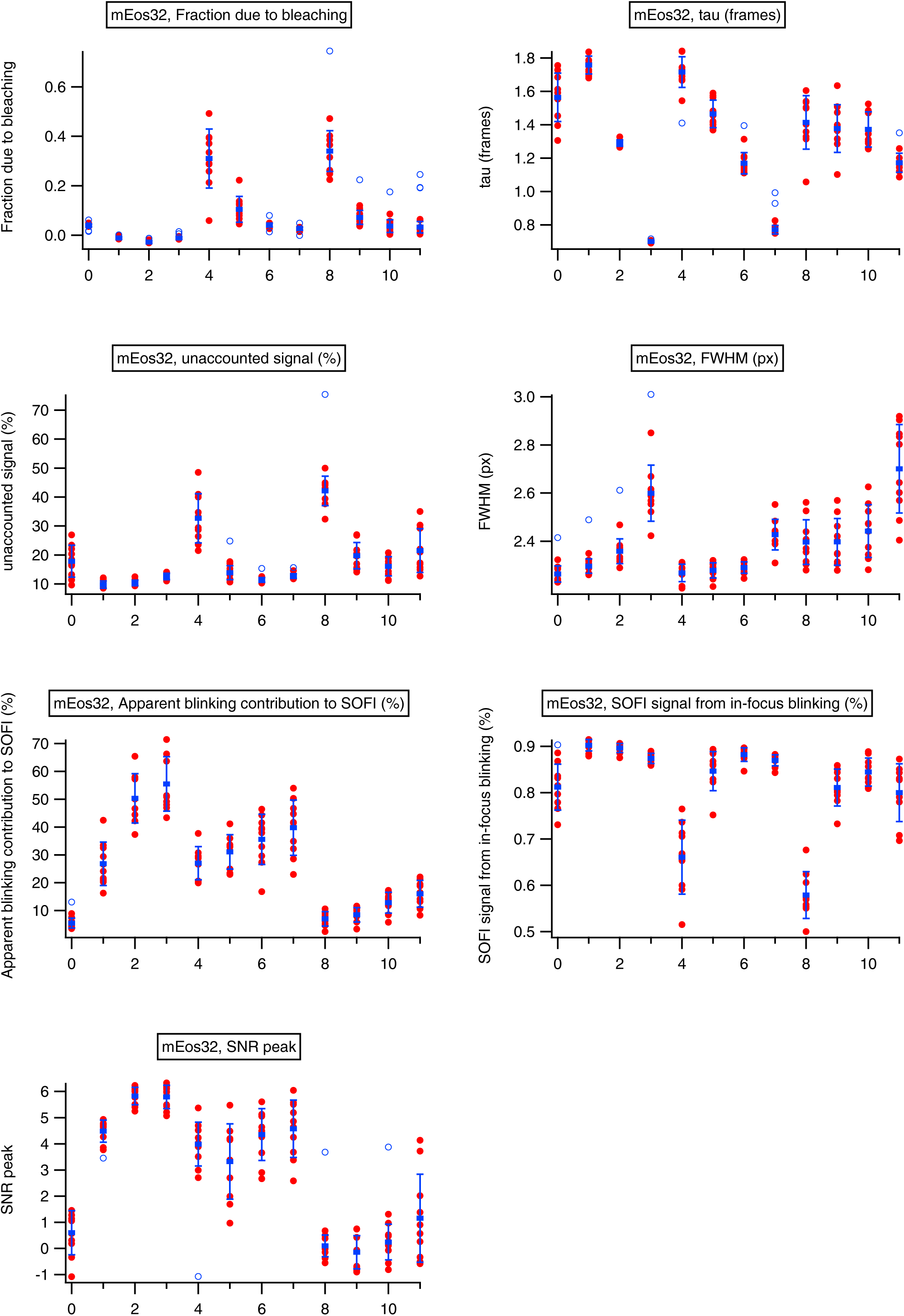
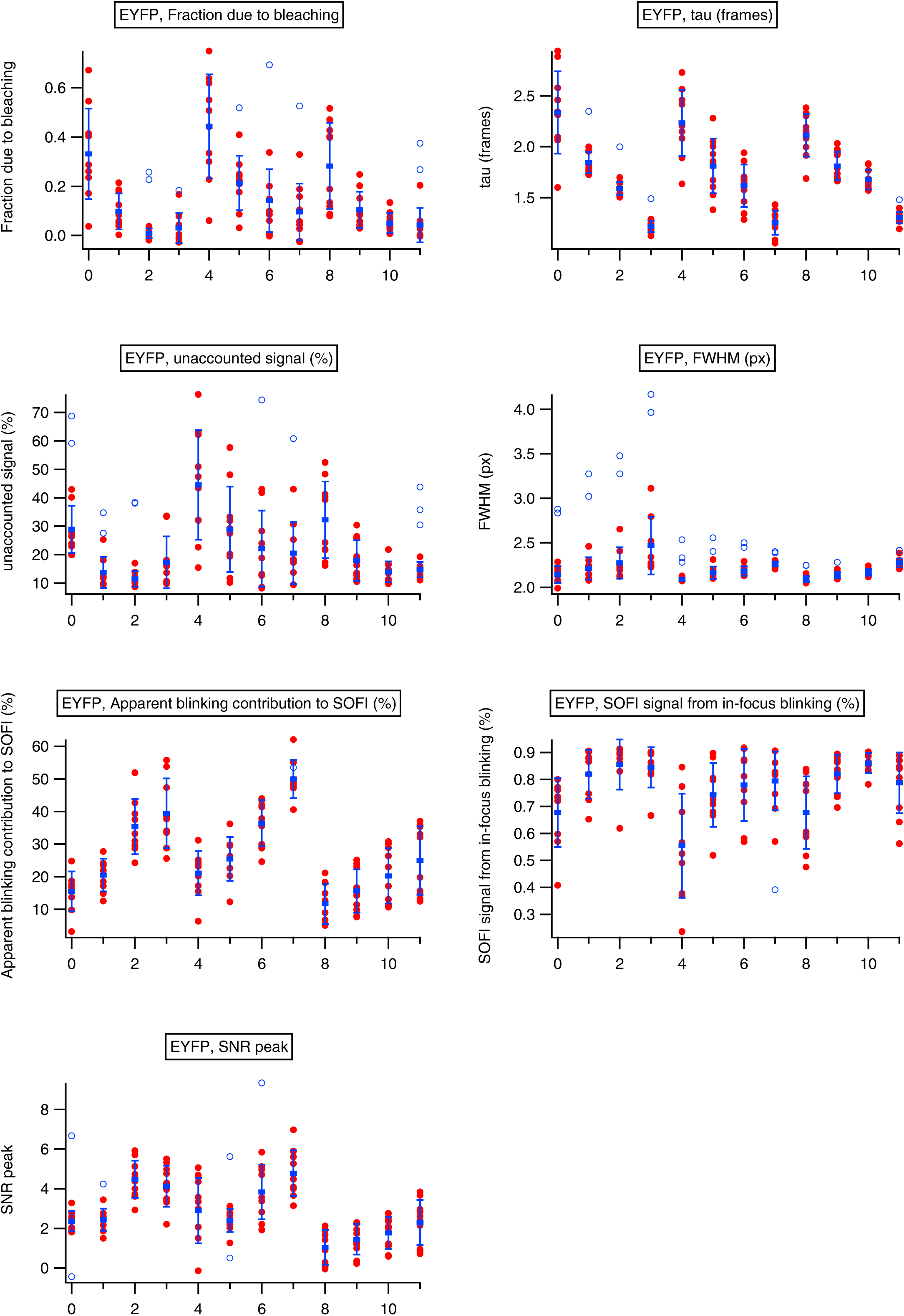
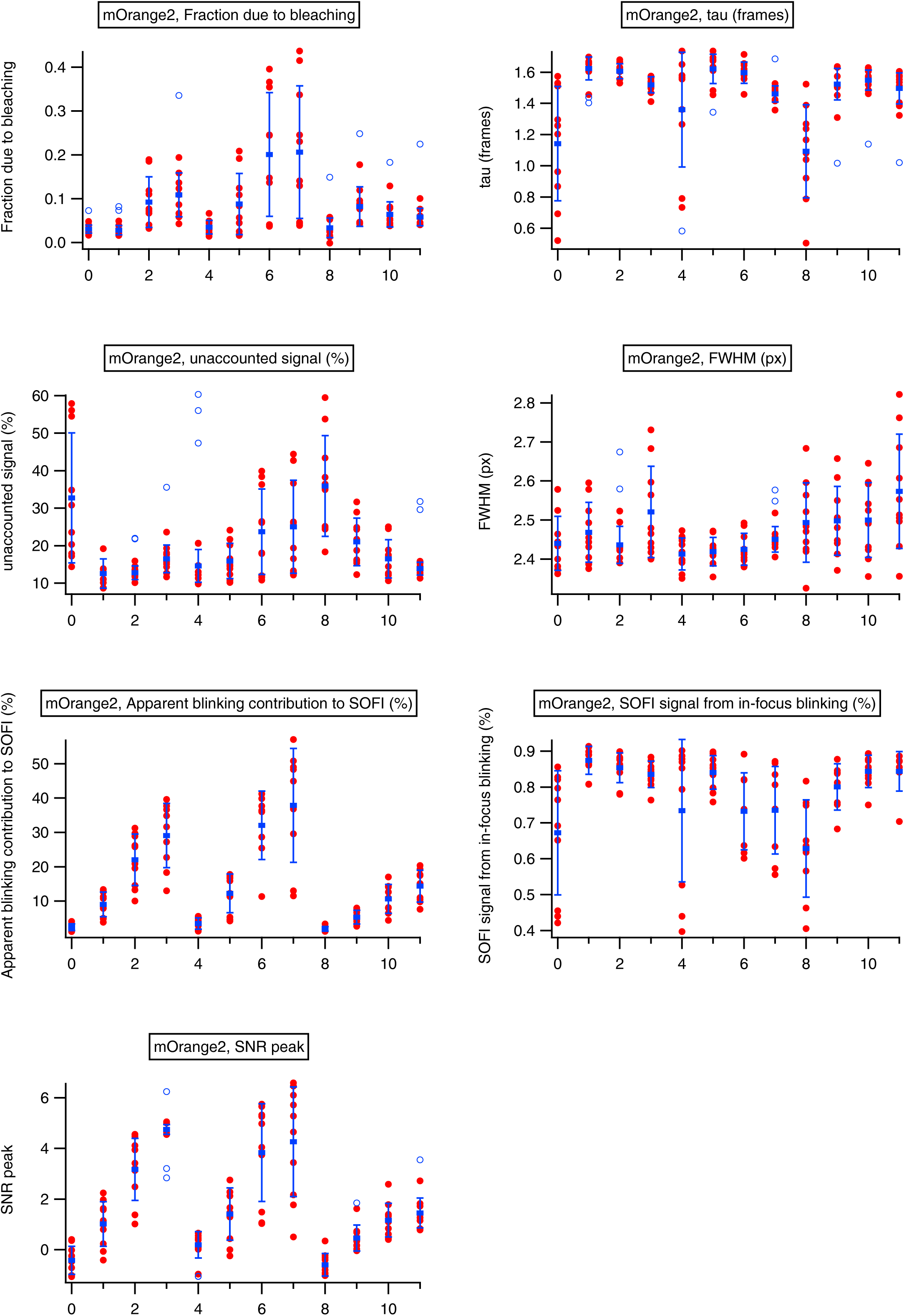
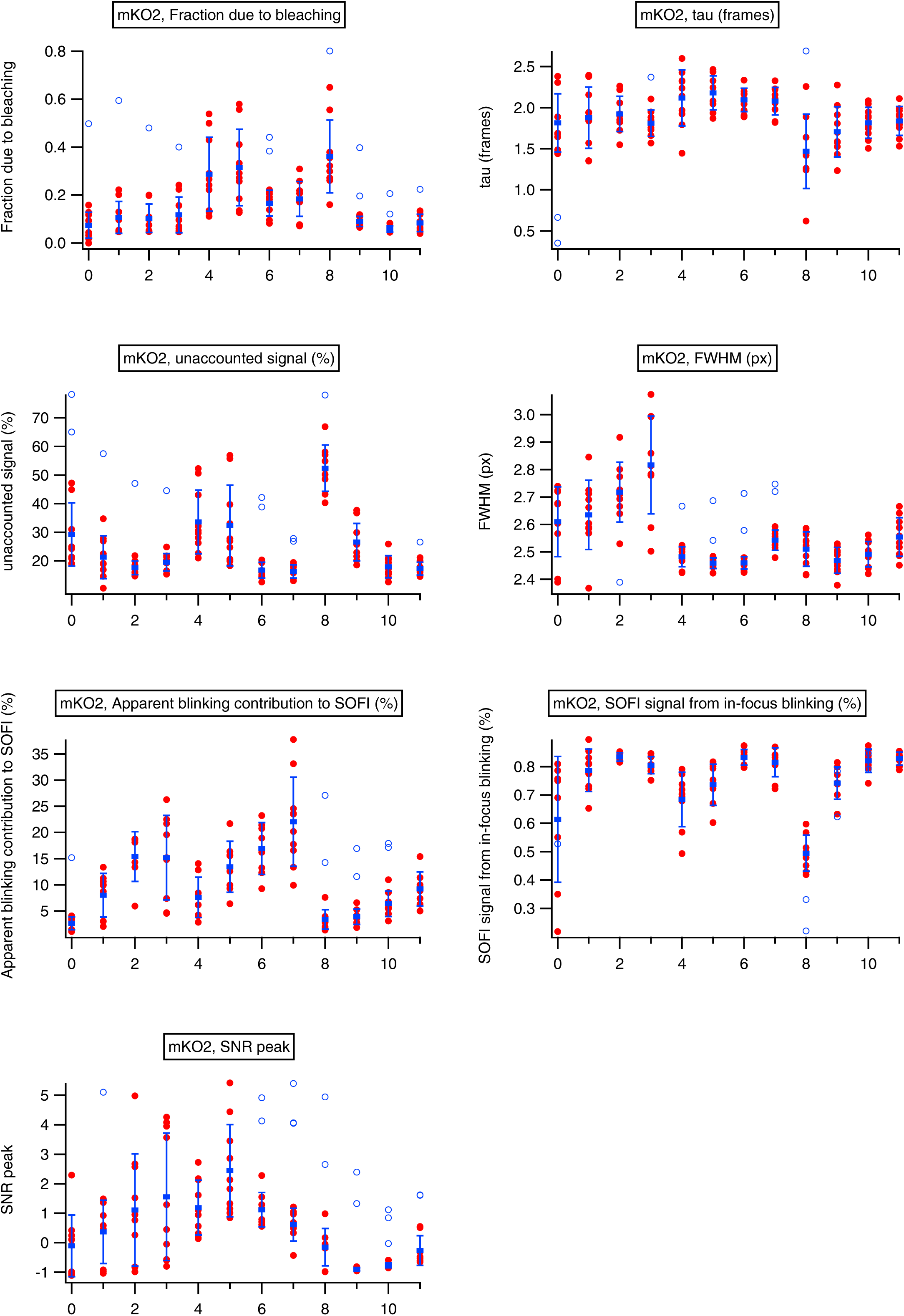
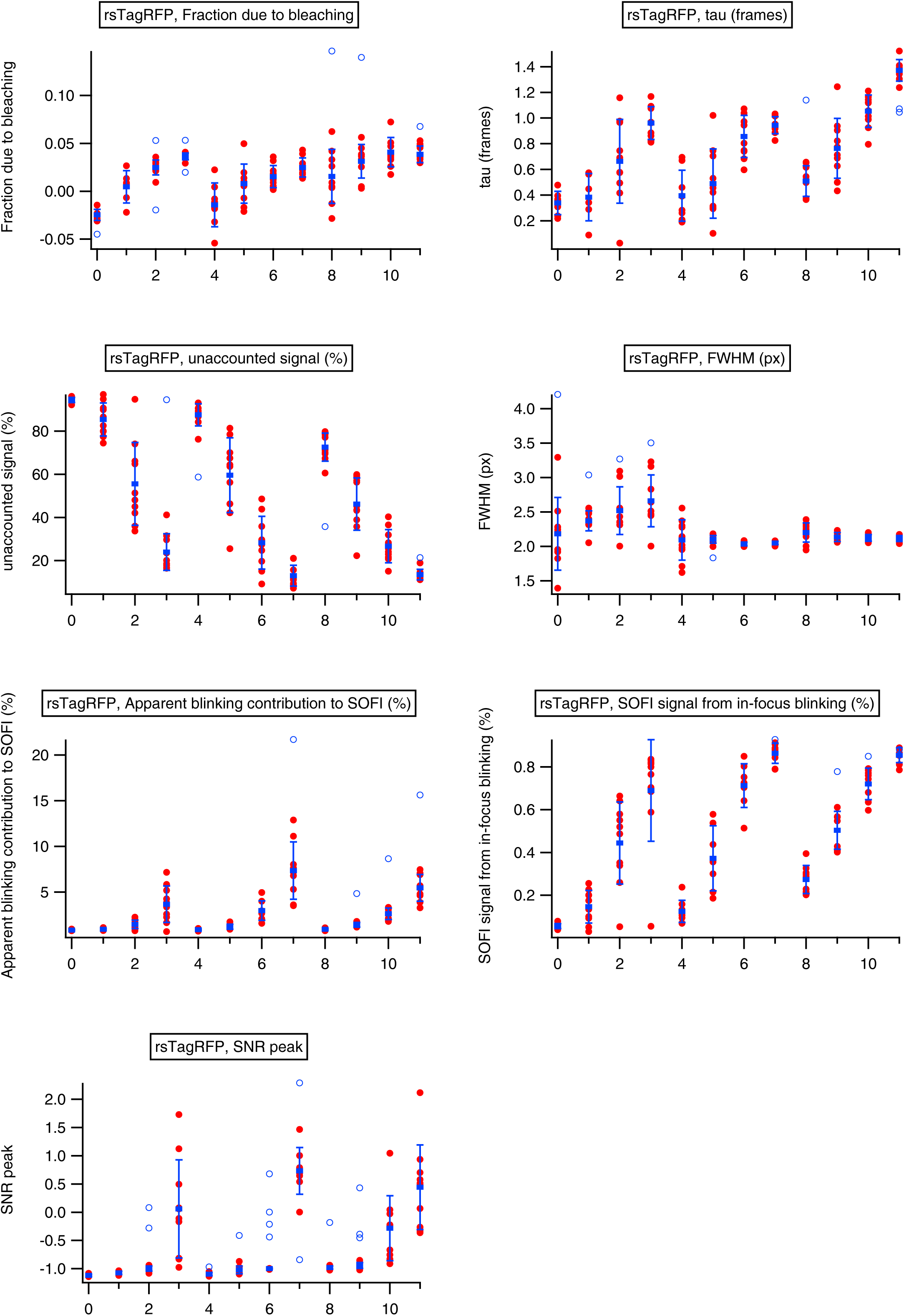
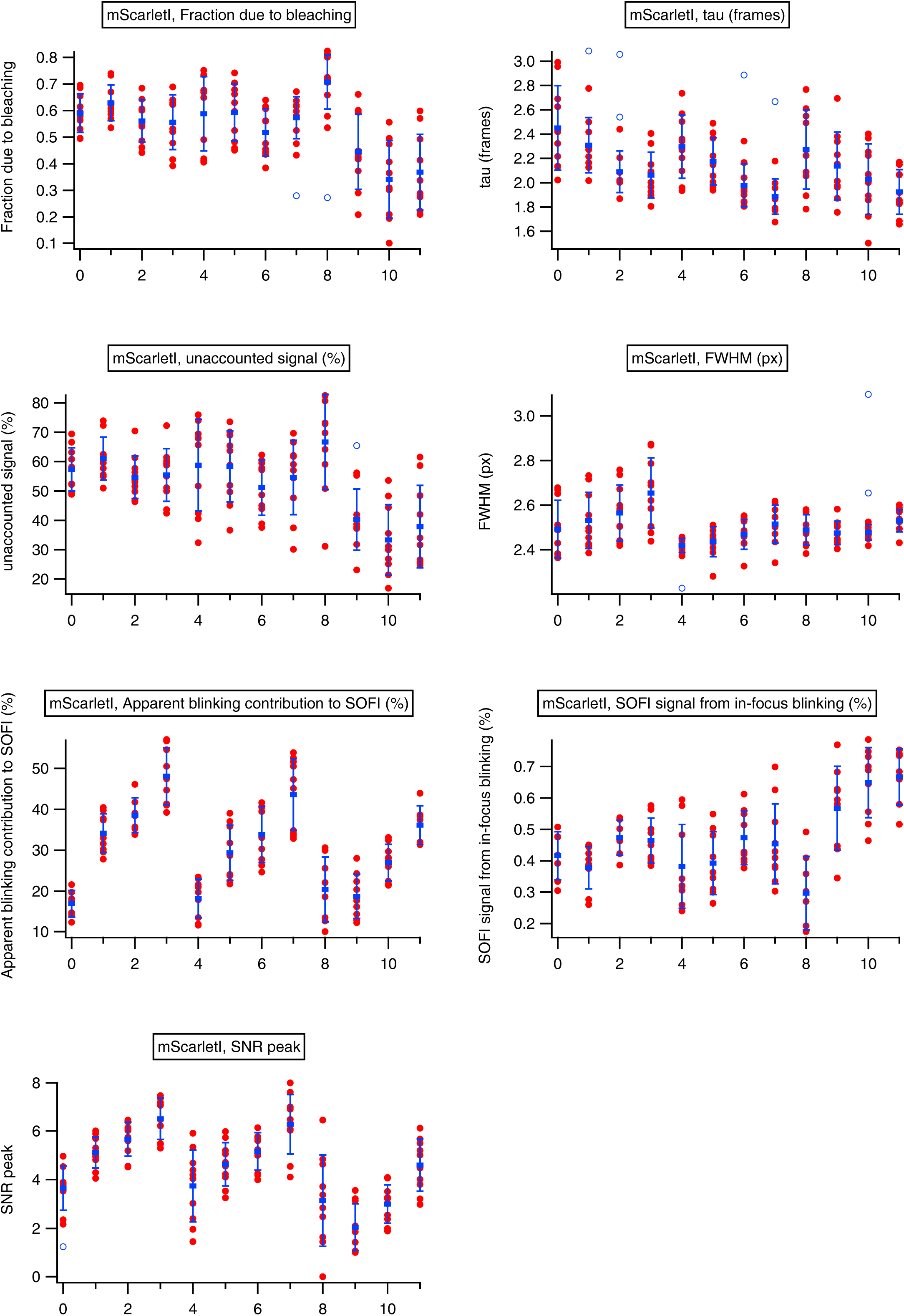
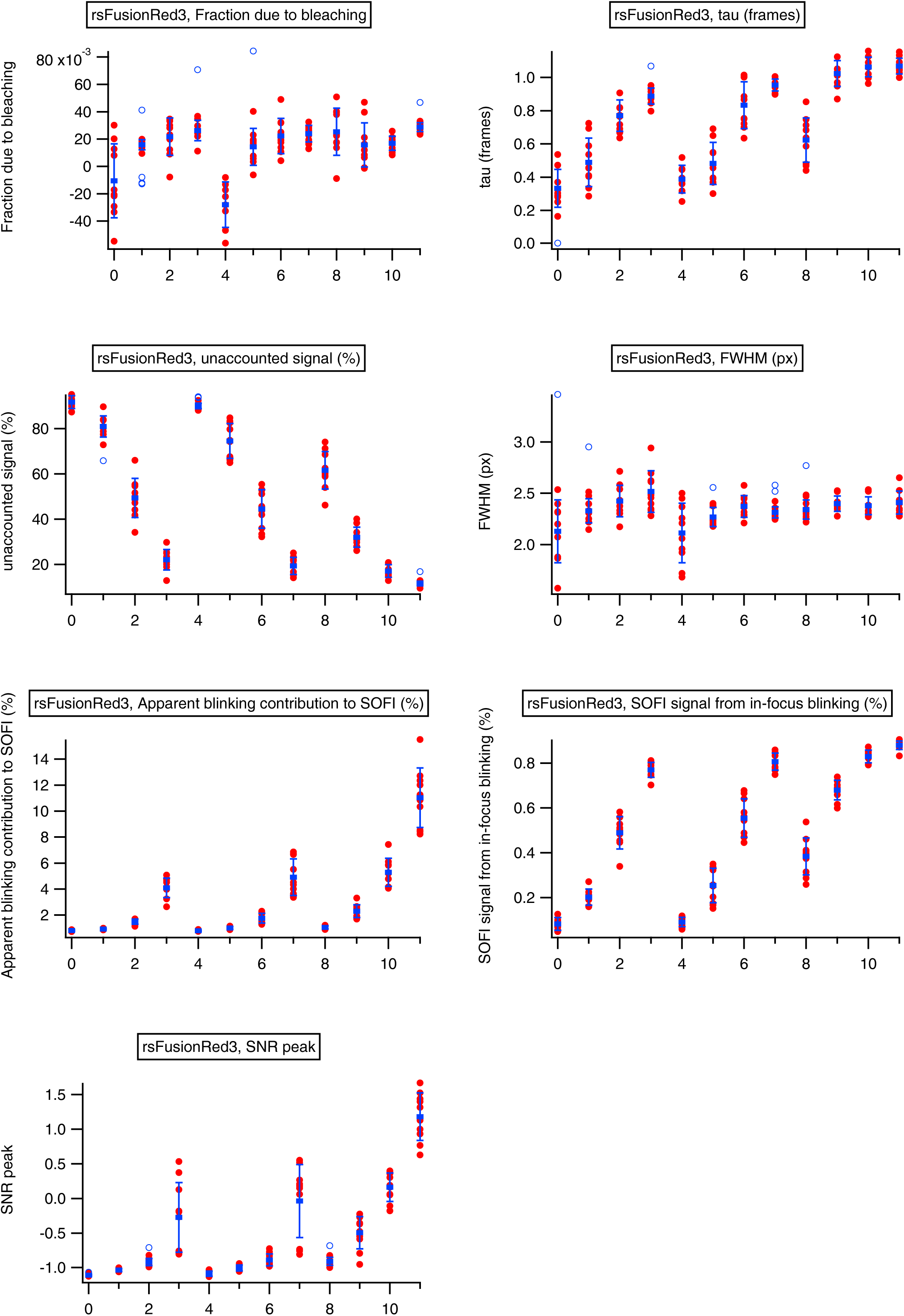
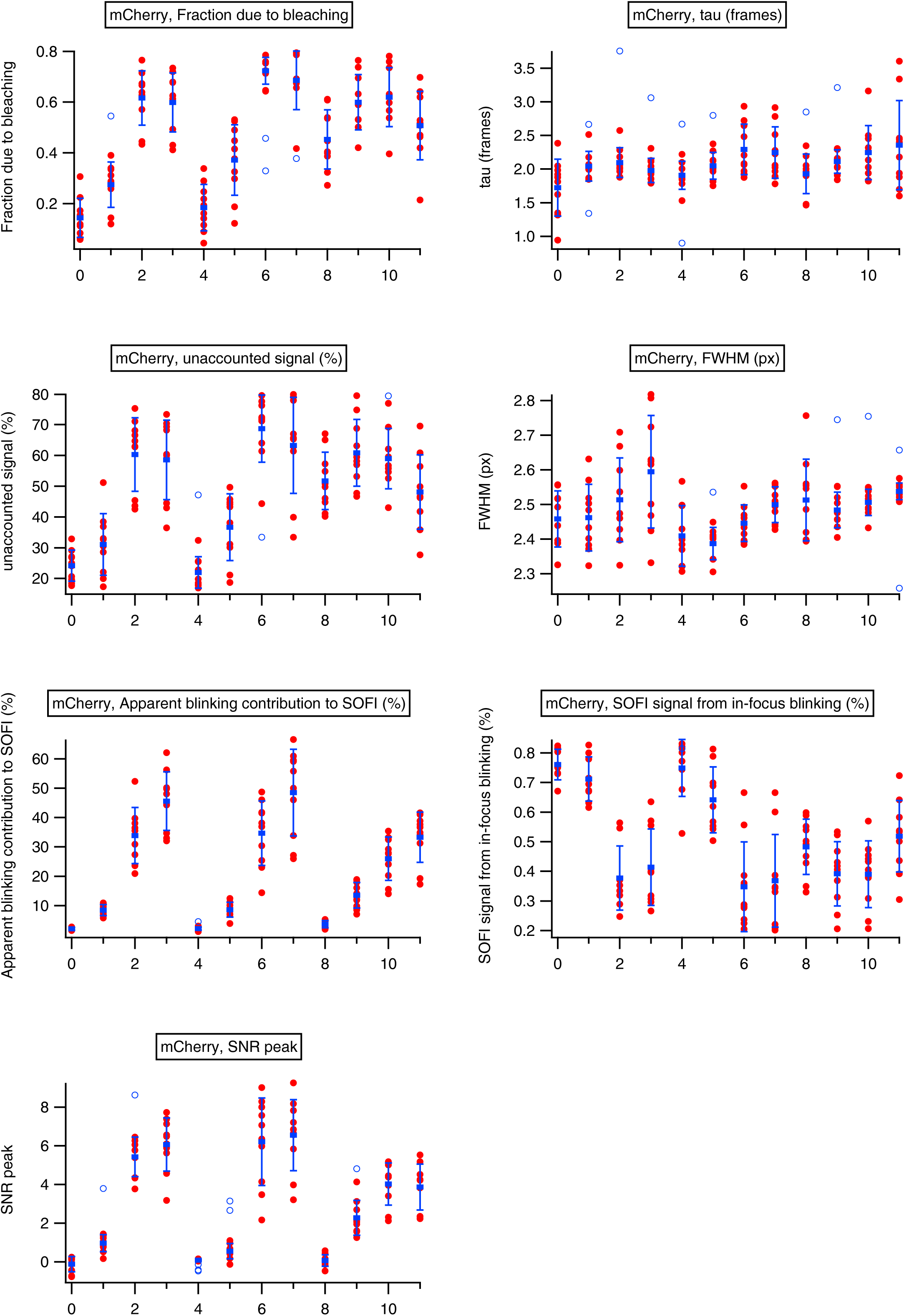

